# Chronic Infection Perturbs the Affinity Hierarchy of Antiviral B Cells

**DOI:** 10.1101/2025.08.21.671513

**Authors:** Mirela Dimitrova, Tiago Abreu-Mota, Jonas Fixemer, Weldy V. Bonilla, Anna-Friederike Marx, Min Lu, Karen Tintignac, Anna Lena Kastner, Yusuf Ismail Ertuna, Marianna Florova, Matias Ciancaglini, Kerstin Narr, Karsten Stauffer, Julien Roux, Philippe Demougin, Ingrid Wagner, Doron Merkler, Daniel D. Pinschewer

## Abstract

The germinal center (GC) subjects antigen-specific B cells to a Darwinian selection process. Whether and how persistent viral infection perturbs the intended affinity hierarchy remains ill-defined.

Here we transferred monoclonal lymphocytic choriomeningitis virus-specific B cells into persistently infected hosts. High affinity B cells expanded vigorously, forming GCs and abundant antibody-secreting cells. When failing to gain the upper hand over the virus, the expanded B cell population contracted, ending in its quasi-complete disappearance, a process termed “attrition”. In stark contrast, low-affinity B cells expanded and persisted irrespective of high viral loads. B cell attrition was associated with phenotypic and transcriptional alterations including a prominent Blimp-1 transcriptional signature in high-affinity GC B cells. Blimp-1-deficient B cells were resistant to attrition, suggesting a B cell-intrinsic process. Moreover, exogenously supplied antibody feedback prevented attrition, indicating the latter resulted from excessive stimulation.

Our findings reveal that in chronic viral infection the incessant activation by overwhelming amounts of antigen perturbs B cell affinity hierarchies by preferentially dysregulating high-affinity B cells.

## Introduction

The World Health Organization estimates that more than 300 million people worldwide live with hepatitis B and/or hepatitis C virus (HBV, HCV)^1,2^, and approximately 38 million people are carriers of HIV ^3^. Despite potentially fatal long-term consequences, a functional cure for HBV or HIV infection remains currently out of reach ^4-7^. Chronic viremic infections tend to subvert adaptive immune defense as exemplified by “T cell exhaustion”, which is intimately linked to persistently high antigen levels ^8-17^. Evidence it accumulating that not only T cells but also humoral immune responses are suppressed in chronic viral infection. This is evident in delayed and inadequate antibody responses to HCV and HIV in humans and to lymphocytic choriomeningitis virus (LCMV) in mice ^18-21^. The B cell compartment in chronic HBV and HIV infection undergoes phenotypic alterations as reflected in the accumulation of atypical memory B cells and plasmablasts, suggesting biased end-differentiation ^22-26^. Additionally, B cells in the peripheral blood of patients exhibit elevated levels of inhibitory receptors such as FcRL4 and PD-1 ^22,23,27,28^, and in the case of HBV infection show impaired differentiation into antibody-secreting cells (ASCs) as well as inadequate immunoglobulin secretion ^27,28^. We and others have reported that type I interferon (IFN-l) responses at the onset of chronic infection can lead to *“decimation”* of virus-specific B cells ^29-31^, reducing antiviral B cell expansion by ∼10-fold ^32^. While contributing to delayed neutralizing antibody (nAb) responses, the aforementioned effect size seems insufficient to fully account for the inadequacy of antibody responses to chronic viral infection, necessitating further investigations into the limitations of B cell responses in this context. B cell-based protection has significant potential as a means to contain persistent viral diseases. Timely nAb responses upon primary as well as secondary HCV infection herald spontaneous immune-mediated viral clearance ^33-36^. Similarly, high frequencies of HIV-specific memory B cells (MBCs) are a frequent characteristic of HIV post-treatment controllers that suppress viral loads for several years after cessation of antiretroviral therapy ^37^. Moreover, potent anti-HIV antibody responses are often associated with HIV elite control ^38-40^ and spontaneous clearance of HBV carriage goes along with seroconversion to protective anti-HBs antibodies ^41^. Prominent glycan shields impede, however, antibody access to neutralizing epitopes on the envelope glycoproteins of HIV, HCV and LCMV, rendering these viruses extremely challenging targets for nAb induction ^42-44^. Even when presented to the immune system in the context of vaccination rather than in chronic infection, neutralizing antibody (nAb) responses are weak or not elicited at all ^21,44-46^. Accordingly, only a minority of HIV-infected individuals succeed after years of viremic infection in mounting broadly neutralizing antibodies (bnAbs) that cover a majority of viral genomic variants ^33,47-52^. With rare exceptions, however, such bnAbs fail to neutralize the donating patients’ autologous virus at the time of antibody cloning ^33,39^. Known limitations curtailing bnAb responses comprise that in HIV-negative individuals only about one naïve precursor of bnAb-producing B cells is found per million B cells ^53^. Moreover, these germline antibodies bind to their target with only low to intermediate affinity and cannot neutralize the virus ^54,55^. Therefore and unlike for acute viral infections such as SARS-CoV-2, influenza A virus or vesicular stomatitis virus ^56-59^, the evolution of HIV-bnAbs necessitates substantial affinity maturation ^54,55^. The continuous maturation of serum antibody affinity over time ^60^ is the end result of somatic B cell receptor (BCR) hypermutation in the germinal center (GC) dark zone, followed by competition of B cells for antigen on follicular dendritic cells and the selection of high-affinity clones by cognate follicular T helper cells in the light zone ^61-70^. Inter-clonal competition of B cells is not, however, limited to the GC. Even very-low affinity clones can be recruited to GCs ^71-73^ but affinity-based pre-GC competition for T help can restrict GC access ^72,74^, a limitation that is relaxed under conditions of abundant cognate T help ^75^. These several layers of Darwinian selection predict an enrichment of high-affinity clones in the GC over time, alongside a quasi-complete elimination of lower-affinity competitors. Single-cell GC repertoire analyses from vaccinated animals have, however, revealed that the average BCR affinity increases over time while the GC reaction as a whole maintains a broad range of affinities throughout the response, notably including low-affinity BCRs ^76-79^. Chronic viral infection is predicted to set exceptional conditions for B cell selection, since antigen is available in excess and the GC response is tasked to convert B cell clones of very low abundance and low starting affinity into a high-affinity bnAb response ^51,52,54,55^. It remains unknown but would be important to understand how these exceptional parameters impact affinity hierarchies in antiviral B cell responses.

Here we report that in chronic viral infection, transient expansion of high-affinity B cells is followed by their Blimp-1-dependent clonal attrition, whereas low-affinity B cells mount sustained responses. These findings indicate that persistently high levels of viral antigen in chronic infection can perturb the affinity hierarchy of responding B cell populations.

## Results

### Abortive expansion of antiviral B cells when transferred at very low numbers into chronically infected hosts

Considering the very low precursor frequency of bnAb-producing B cells in humans ^53^ and the challenges related to the efficient recruitment of such low-frequency B cell populations into GC responses to vaccination ^80^ we set out to study how very low numbers of nAb-producing B cells respond to chronic viral infection in mice. HkiL donor B cells are engineered to express the KL25 antibody, which binds the LCMV-WE strain envelope glycoprotein (GP) at 5 nM affinity and exhibits potent virus-neutralizing activity ^81^. Consistent with earlier reports ^80,82,83^ we found that the splenic take of adoptively transferred HkiL cells was in the range of ≤5% (Fig. S1). Taking this uptake rate as a basis of calculation we engrafted either 10, 100 or 1000 HkiL cells in the spleen of mice that underwent chronic infection with an engineered Clone 13-based LCMV strain expressing the WE strain GP ^84^ (rCl13/WE; Fig. 1A). Adoptive B cell transfer (Tf) was conducted six days after virus administration to avoid IFN-I-induced decimation in the first few days after LCMV infection ^29-31^ and to mimic the continuous recruitment of antiviral antibody-producing B cell clones into the response to ongoing HIV-infection ^51^. Four weeks after engraftment of 100 or 1000 HkiL cells their expanded progeny populations (CD45.1^+^) were readily detected in the respective recipients (100-cell recipients, 1000-cell-recipients) and displayed mostly a GL7^+^CD38^-^CD138^-^ GC phenotype (Fig. 1C,D, Fig. S1A). In contrast recipients of 10 cells (10-cell-recipients) did not harbor HkiL progeny at numbers clearly exceeding technical backgrounds of mice without adoptively transferred HkiL cells (“noTf”). In keeping therewith, 100-cell-recipients and 1000-cell-recipients but not 10-cell-recipients suppressed viremia to below detection limits at week 4 (Fig. 1E). The serum of 10-cell-recipients did not contain any detectable KL25 antibody at week 4 either, but the same was clearly detected at week 2, suggesting that HkiL cells mounted a transient antiviral response (Fig. 1B).

**Figure 1:**
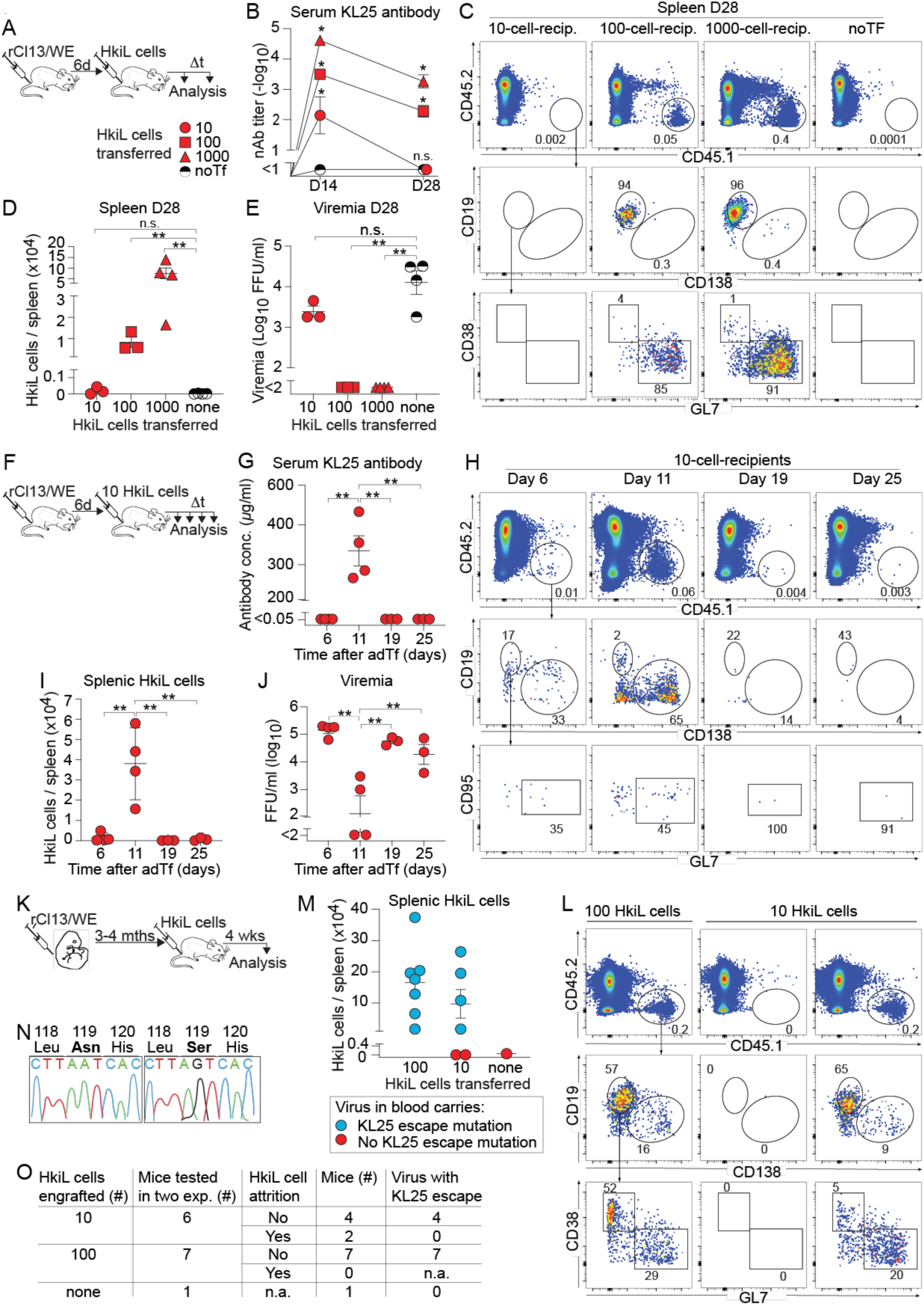
Abortive expansion of antiviral B cells when transferred at very low numbers into chronically infected hosts. A-E: We infected mice with rCl13/WE on d-6. On d0 we engrafted either 10, 100 or 1000 HkiL cells or none (noTF), collected serum over time and analyzed HkiL progeny in spleen on d28 (A). Neutralizing antibody titers over time (B) and representative FACS plots from spleen on d28 (C; 3 mice in 10- and 100-cell groups, 4 mice in the remainder groups) pre-gated on live lymphocytes (Fig. S1A) are shown. HkiL progeny (CD45.1^+^, top) were analyzed for expression of CD19 and CD138 (center), and their CD19^+^CD138^—^ subset was analyzed for GL7 and CD38 expression (bottom). Total CD45.1^+^ KL25 progeny counts (D) and viremia (E) were determined on d28. F-I: We infected mice with rCl13/WE on d-6. On d0 we engrafted 10 HkiL cells. Serum was collected and groups of mice were sacrificed on the indicated days (n=4 on d6 and d11; n=3 on d19 and d25). KL25 antibody concentration was determined by ELISA (G) and CD45.1^+^ HkiL progeny (H) were analyzed as in (C) and enumerated (I). Viremia (J) was determined at the indicated time points. K-O: In two separate experiments we infected newborn mice with rCl13/WE. When they were 14-16 weeks old we engrafted them with either 10 or 100 HkiL cells and analyzed them 3-4 weeks later (K; 21 days and 31 days in the two respective experiments). Representative FACS plots (L; n=7 for 100-cell group, n=6 for 10-cell group) and total cell count (M) of splenic HkiL cell progeny. In (M), mice that harbored viruses with KL25 escape mutation are shown in blue, mice with viruses devoid of such mutations are displayed in red. Exemplary fluorograms of Sanger sequencing reactions from viruses in mouse blood (N). The sequence around Aa119 of the viral GP (N) and a summary table of experimental outcome in individual mice (O) are shown (see also Tbl. SI). Numbers in FACS plots indicate the percentage of gated cells (mean of the group). Symbols in (D,E,G,I,J,M) show individual mice with the mean±SEM indicated, in (B) they show the mean±SEM (n=3-4 per group). Data in (A-J) show one out of two similar experiments. Panels (K-O) report combined results from two experiments. Two-way ANOVA with Tukey’s post-test was performed in (B) reporting for each time point significant differences in comparison to the “noTf” group. 1-way ANOVA with Dunnett’s post-test was performed in (D,E), comparing to the noTF group. In (G,I,J) we performed 1-way ANOVA with Tukey’s post-test, comparing the means of all groups and showing only significant differences. *: p<0.05; **: p<0.01; n.s.: not statistically significant.

Accordingly, a time course analysis in the spleen of 10-cell-recipients showed that HkiL cells expanded until day 11 after engraftment, forming a substantial population of predominantly ASC-differentiated (CD138^+^B220^low/int^) progeny, but contracted to background levels by day 19 and thereafter (Fig. 1H,I). This initial expansion and subsequent collapse of the HkiL cell population was reflected in a transient KL25 antibody response in serum and a concomitant suppression of viremia on day 11, which rebounded by day 19 (Fig. 1G,J). To extend and generalize our observations we performed HkiL transfer experiments in neonatally infected carriers of rCl13/WE, which exhibit life-long unchecked viremia at levels that are equivalent to or higher than those observed when chronic infection is established in adult life ^85,86^. Neonatally infected carriers develop, however, antiviral CD8 T cell tolerance^87^ and are virtually devoid of the antiviral inflammatory reaction^88,89^, which in adult infected animals can profoundly impact antiviral B cell responses^29-31^. When engrafting 100 HkiL cells into rCl13 carriers, all animals exhibited a clearly detectable population of HkiL cell progeny four weeks later, most of which were GC B cells (Fig. 1K,L,M). Interestingly, analogous populations were also found in four out of six carriers engrafted with 10 HkiL cells in two independent experiments, whereas one animal in each experiment was devoid of detectable HkiL cell populations. We found that in all 10-cell- and 100-cell-recipients with detectable HkiL cell populations the virus persisted but had acquired typical KL25 escape mutations at amino acid 119 of the glycoprotein ^81,90,91^ (Fig. 1M, blue symbols; Fig. 1N,O; Tbl. SI). No such mutations were found in animals that were devoid of detectable HkiL cell progeny (Fig. 1M, red symbols; Fig. 1N,O; Tbl. SI). Altogether these observations suggested a dichotomous development of adoptively transferred HkiL cells in chronic LCMV infection. When transferred in higher numbers, these antiviral B cells gained the upper hand i.e. eliminated the virus in adult rCl13/WE infection or forced it into mutational escape in neonatally infected carriers. When engrafted at critically low cell numbers, however, the transferred HkiL cells were subdue after initial expansion, a process henceforth referred to as “attrition”.

### Chronic infection with high-affinity but not low-affinity virus antigen causes B cell attrition

The observation that HkiL cells persisted in neonatally infected carrier mice whenever the virus underwent mutational escape was intriguing, prompting us to study the response to N119S-mutant virus in the adult infection setting. These attempts were, however, encumbered by the inability of rCl13/WE to persist when more than 10 HkiL cells were engrafted. Hence, we established a reverse genetic system for the LCMV strain Docile (DOC)^92,93^, a variant of the WE strain which establishes more robust and long-lasting viremia than rCl13/WE (Fig. S2A,B). We generated a pair of viruses, DOC (wildtype virus) and a KL25 low-affinity variant (DOC-LAV). DOC-LAV differed from DOC by only one single point mutation in the viral glycoprotein (N119S), an escape mutation without associated fitness cost that was frequently detected in the experiments to Fig. 1K-O (Tbl.I) and is known to reduce KL25 binding by ≥1000-fold ^81,94^. Mice were infected with either DOC or DOC-LAV and were engrafted six days later with either 10, 30 or 100 HkiL cells (Fig. 2A). When analyzing spleens of DOC-infected mice 24 days later, HkiL cell progeny were readily detected in 30- and 100-cell-recipients but had seemingly undergone attrition in 10-cell-recipients (Fig. 2B,C, S2C). In stark contrast, a sizeable population of HkiL cells persisted in DOC-LAV-infected 10-cell-recipients and consisted mostly of GC B cells (Fig. 2B,C, S2C). KL25 antibody production was readily detected in all groups of mice on day 8 and on day 15 after HkiL cell engraftment, but in DOC-infected 10-cell-recipients dropped to below detection limits on day 24 (Fig. 2E). Analogously to these findings in the spleen, HkiL cells persisted in lymph nodes of DOC-LAV-infected but not DOC-infected 10-cell-recipients (Fig. S2F,G), whereas HkiL cell progeny in the bone marrow were not clearly over noTf background, irrespectively of the infecting virus (Fig. S2D,E). As hoped for and unlike in adult rCl13/WE infection, viremia persisted in all groups of mice throughout the observation period (Fig. 2D, compare Fig. 1E). Still, HkiL cells afforded partial suppression of DOC viremia, whereas replication of the KL25 escape variant DOC-LAV was unaffected, as expected (Fig. 2D)^81^. Sequence analysis of the persisting DOC viruses on day 24 after HkiL cell transfer evidenced an unmodified DOC GP sequence in 10-cell-recipients but mutational escape in 30-cell- and 100-cell-recipients, correlating with the attrition and persistence of HkiL cells, respectively (Fig. 2F,G; Tbl. SI). These analyses were complemented by immunofluorescence stains of spleen sections from DOC and DOC-LAV-infected 10-cell-recipients on day 24 after engraftment. The total number of GCs per section were comparable in the two groups (Fig. 2H,I,J), yet HkiL cell progeny (CD45.1^+^) were consistently found in PNA^+^ GC structures of DOC-LAV-infected 10-cell-recipients but were virtually absent from GCs of DOC-infected 10-cell-recipients (Fig. 2H,I,K). These observations on clonal attrition of high-but not low-affinity B cells violated the principles of Darwinian selection and contrasted with earlier studies in the context of protein vaccination, showing that high-affinity B cells mounted equal if not stronger responses than their low-affinity counterpart ^73,95^.

**Figure 2:**
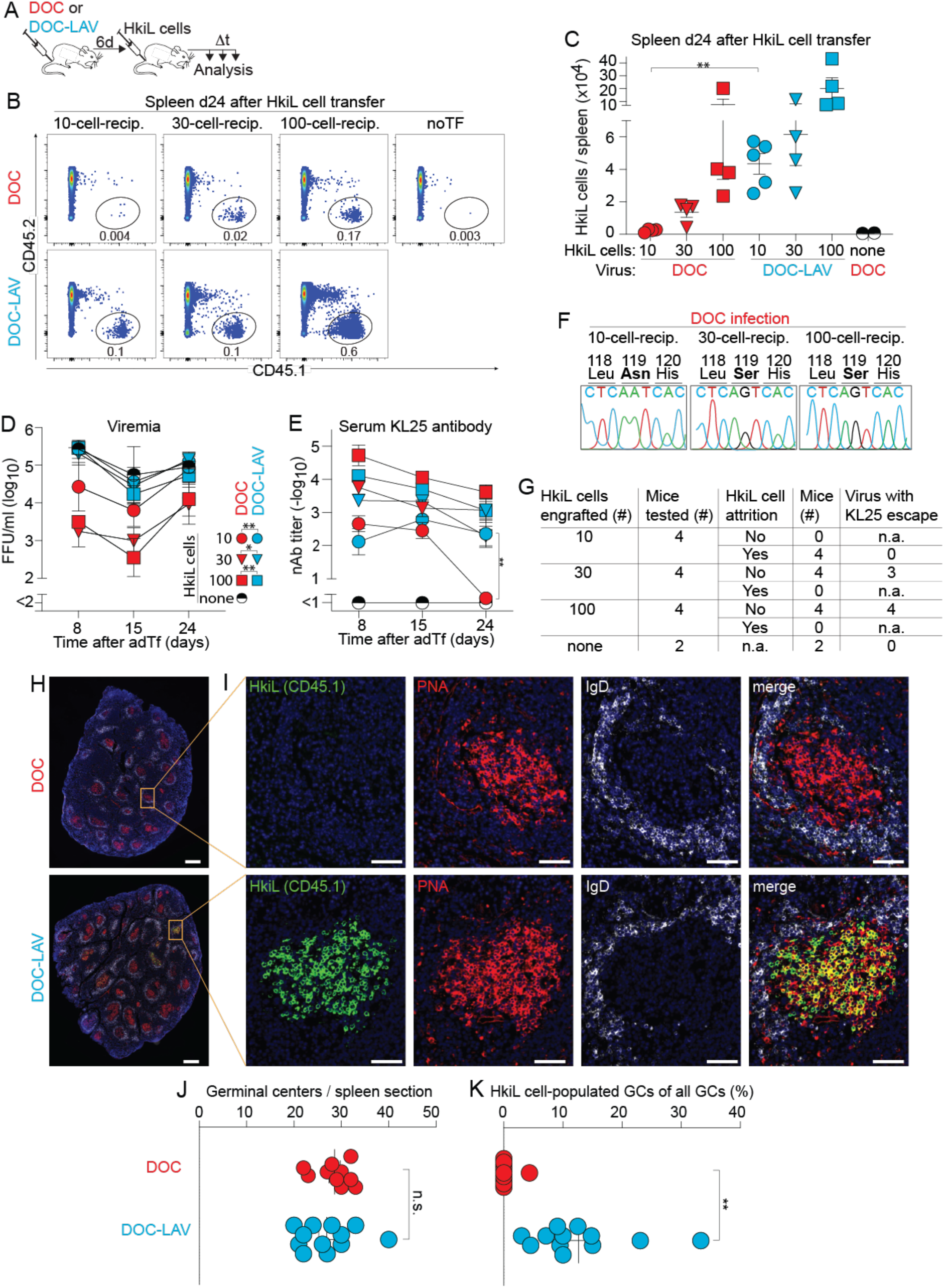
Chronic infection with high-affinity but not low-affinity viral antigen causes B cell attrition. We infected mice with DOC or DOC-LAV on d-6. On d0 we engrafted either 10, 30 or 100 HkiL cells or none (noTF), collected blood and serum samples over time and analyzed HkiL cell progeny in spleen on d24 (A-C). Representative FACS plots (B; n=4 mice per group except DOC-LAV-infected 10-cell-recipients (n=5) and noTf (n=2)) pre-gated on live lymphocytes (Fig. S1A) show HkiL progeny (CD45.1^+^). The cells’ expression of B220 and CD138, and the GL7 / CD38 profile of the B220^+^CD138^—^ subset are reported in Fig. S2C. HkiL progeny were enumerated (C) and viremia (D) as well as nAb titers (E) were determined over time. Exemplary fluorograms of Sanger sequencing reactions of viruses persisting in DOC-infected 10-cell-, 30-cell- and 100-cell-recipients on d24 (F). The sequence around Aa119 of the viral GP and a summary table of experimental outcome in individual mice (G) are shown (see also Tbl. SI). Spleen sections were stained for CD45.1 (HkiL cells; green), PNA (GC; red), IgD (white) and DAPI (H,I). The total number of GCs per section (J) and the percentage of GCs containing three or more CD45.1^+^ cells (K) were determined. Numbers in FACS plots indicate the percentage of gated cells (group mean; B). Symbols in (C) show individual mice. Symbols in (D,E) represent the mean±SEM of n=4-5 mice per group (except noTf group: n=2). In (H,I) representative images from 10 spleen sections of 4-5 mice per group are shown, symbols in (J,K) represent individual spleen sections (up to three sections per mouse taken at ∼100 µm distance from each other) with the mean±SEM indicated. Magnification bars: 500 µm (H) and 50 µm (I). One-way ANOVA with Tukey’s post-test of log-converted values was performed in (C), and significant differences between groups receiving the same number of HkiL cells are reported. Repeated measures ANOVA was performed for a pairwise comparison of DOC- and DOC-LAV-infected groups receiving the same respective number of HkiL cells (D). Two-way ANOVA with Tukey’s post-test was performed in (E) and the only significant difference between groups receiving the same respective number of cells is reported for the respective time point. Unpaired Student t-tests were performed in (J,K). Only statistically significant differences are shown in (C-E). Panels (B-F) show one representative out of two similar experiments. *: p<0.05; **: p<0.01; n.s.: not statistically significant.

### B cell attrition is due to excessive amounts of viral antigen and occurs independently of IFN-I signaling

To test the hypothesis that attrition was inherently linked to persistent infection, we tested the response of HkiL cells to acute infection and vectored vaccination. We infected mice with either one of two engineered variants of the non-persisting LCMV strain Armstrong, expressing either WE-GP or its low-affinity N119S-mutant (rARM; rARM-LAV), engrafted them with 10 HkiL cells and analyzed these cells’ progeny twenty days later (Fig. 3A). Comparable numbers of HkiL progeny were found in both infection settings, indicating that attrition was not observed in the context of acute, non-persisting infection (Fig. 3B,C, S3A). To corroborate and extend these conclusions we engineered Pichinde virus, which replicates poorly in mice (Fig. S3B) but shows utility as a viral vaccine vector ^96^, to express either WE-GP or the low-affinity N119S variant (rPICV; rPICV-LAV) instead of its own glycoprotein. When infected with rPICV, 10-cell-recipients formed a substantial population of HkiL cell progeny, whereas in rPICV-LAV-infected recipients HkiL cell progeny were significantly less abundant (Fig. 3D-F, S3C). These observations demonstrated that 10 engrafted HkiL cells were perfectly able to mount sustainable responses to high-affinity WE-GP provided the latter was presented in the context of acute infection or vaccination.

**Figure 3:**
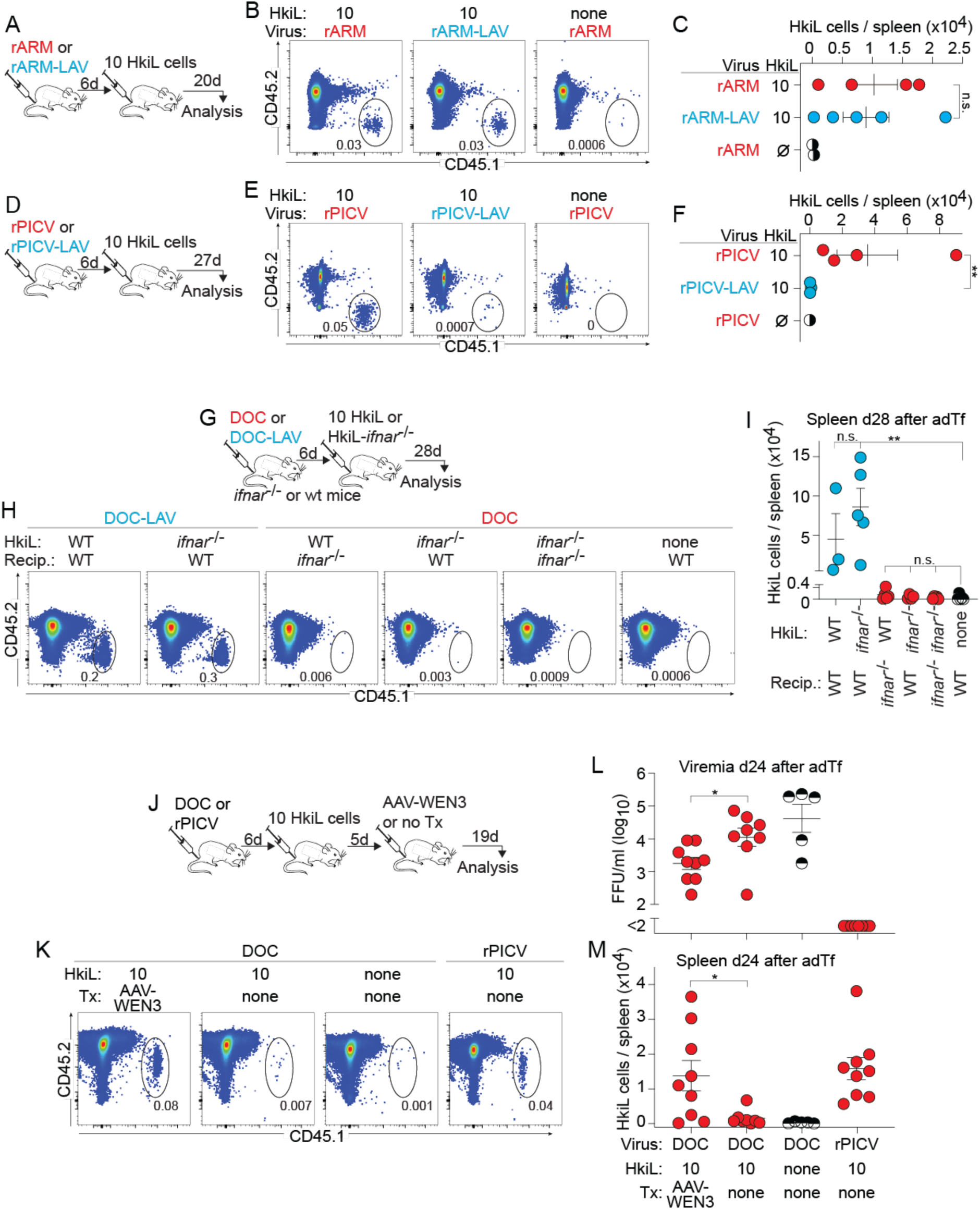
B cell attrition is due to excessive amounts of viral antigen and occurs independently of IFN-I signaling. A-C: We infected mice with rARM or rARM-LAV on d-6 and on d0 engrafted 10 HkiL cells or none (noTf) (A). HkiL progeny in spleen were analyzed and enumerated on d20 (B,C). Representative FACS plots (B; n=4 mice except n=2 mice in noTf group) pre-gated on live lymphocyte (Fig. S1A). The cells’ expression of B220 and CD138, and the GL7/CD38 profile of the B220^+^CD138^—^ subset are reported in Fig. S3A. D-F: We infected mice with rPIC or rPIC-LAV on d-6 and on d0 engrafted 10 HkiL cells or none (noTF) (D). HkiL progeny in spleen were analyzed and enumerated on d27 (E,F). Representative FACS plots (E; n=4 mice for rPIC, n=3 mice for rPICV-LAV group, n=1 for noTF) pre-gated on live lymphocyte (Fig. S1A). The cells’ expression of B220 and CD138, and the GL7/CD38 profile of the B220^+^CD138^—^ subset are reported in Fig. S3C. G-I: We infected *ifnar*^-/-^ and WT control recipients with DOC or DOC-LAV on d-6. On d0 we engrafted them with either 10 HkiL cells or 10 HkiL-*ifnar*^-/-^ cells or left them without cell transfer (”none”, noTF) in the combinations indicated in (H,I). HkiL and HkiL-*ifnar*^-/-^ progeny (CD45.1^+^) were analyzed on d28 (G,I). Representative FACS plots (n=5 except HkiL into DOC-LAV-infected WT mice (n=3) and HkiL-*ifnar*^-/-^ into DOC-infected wt mice (n=4)), pre-gated on live lymphocyte are shown in (H) and CD45.1^+^ progeny were enumerated (I). The cells’ expression of CD19 and CD138, and the GL7/CD38 profile of the CD19^+^CD138^—^ subset are reported in Fig. S3D. J-M: We infected mice with DOC or rPIC on d-6, and on d0 engrafted 10 HkiL cells or none (noTF) as indicated in the chart to (M). On d5 we administered AAV-WEN3 to one DOC-infected group. HkiL progeny in spleen were analyzed on d24 (J). Representative FACS plots (K; n=9 AAV-WEN3-treated HkiL recipients, n=8 recipients without AAV-WEN3, n=9 rPICV-immunized HkiL recipients, n=5 noTF controls) pre-gated on live lymphocytes. Viremia (L) and total splenic HkiL progeny (M) on d24. The cells’ expression of CD19 and CD138, and the GL7/CD38 profile of the CD19^+^CD138^—^ subset are reported in Fig. S3F. Numbers in FACS plots indicate the percentage of gated cells (B,E,H,K) as mean of the group. Symbols in (C,F,I,L,M) show individual mice with the mean±SEM indicated. Student’s t tests of log-converted values were performed in (C,F). Values in (I) were log-converted and analyzed by 1-way ANOVA with Dunnett’s post-test to identify significant differences from the noTF group. Student’s t tests were performed in (L,M). Data in (A-I) show one out of two similar experiments, (L,M) show combined data from two experiments. *: p<0.05; **: p<0.01; n.s.: not statistically significant.

Attrition in chronically infected mice was observed at around two weeks after HkiL cell transfer into pre-infected recipients (compare Fig. 1F-J and below), which differs profoundly from “decimation”. The latter occurs concomitantly with the peak of the IFN-I response during the first three days after LCMV infection and is largely avoided when transferring specific B cells a few days later, as it was done in the present set of experiments ^29-32^. Nevertheless, we set out to formally address the possibility that attrition was due to persisting low-level IFN-I signaling in chronic LCMV infection ^97^. IFN-I receptor-sufficient and -deficient HkiL cells (HkiL; HkiL-*ifnar*^-/-^) expanded comparably when engrafted in DOC-LAV-infected recipients (Fig. 3G-I, S3D), allowing us to test whether B cell-intrinsic and/or -extrinsic IFN-I signaling might contribute to B cell attrition in the context of DOC infection. By transferring either 10 HkiL-*ifnar*^-/-^ cells into wt recipients, 10 HkiL cells into *ifnar*^-/-^ recipients or 10 HkiL-*ifnar*^-/-^ cells into *ifnar*^-/-^ recipients we found that neither B cell-intrinsic IFNAR deficiency, nor recipient IFNAR deficiency or a combination thereof prevented attrition.

We hypothesized that excessive BCR signaling owing to incessant exposure to copious amounts of cognate high-affinity antigen drives B cell attrition in chronic infection. To address this possibility we sought to reduce the antigenic stimulation of HkiL cells by masking their cognate antigen by so-called antibody feedback ^98-100^. Five days after HkiL cell engraftment we treated DOC-infected 10-cell-recipients intramuscularly with an adeno-associated viral vector, a somatic gene therapy, for the continuous high-level production of the DOC-neutralizing antibody WEN3 (AAV-WEN3; Fig. 3J)^81,90,101^. WEN3 competes with KL25 for the same epitope^102^ but typical KL25 escape mutations in the viral GP such as N119S in DOC-LAV do not affect WEN3 binding or neutralization^81,103^. When analyzed 24 days after HkiL cell transfer, AAV-WEN3 treated 10-cell-recipients exhibited significantly lower viral loads than untreated controls (Fig. 3L). Unlike in the latter, where HkiL cells had undergone attrition, HkiL progeny populations were preserved in the majority of AAV-WEN3-treated mice, reaching levels comparable to those of an rPICV-vaccinated reference group (Fig. 3K,M). These results corroborated that the long-term exposure to very high amounts of high-affinity antigen rather than IFN-I signaling accounted for B cell attrition in chronic viral infection.

### HkiL cell populations destined to attrition mount a transient ASC burst and exhibit an altered GC phenotype

To investigate the cellular differentiation processes associated with attrition we compared HkiL cell populations in DOC- and DOC-LAV-infected 10-cell-recipients over time (Fig. 4A). By day 8 after engraftment, HkiL cells responding to DOC infection had given rise to a massive burst of CD138^+^B220^low/int^ ASCs, which contracted by day 14 and reached detection limits on day 31 (Fig. 4B,D). Serum KL25 antibody titers followed these kinetics (Fig. 4E). In contrast, B220^hi^ B cell phenotype HkiL progeny in the same recipients increased in numbers between day 8 and day 14, exhibited predominantly a GC phenotype at both time points and disappeared thereafter (Fig. 4C). These kinetics differed from HkiL cells responding to DOC-LAV infection. B cell phenotype HkiL progeny numbers increased continuously from day 8 to day 31, whereas ASC numbers and antibody titers reached their maximum on day 14 and remained constant thereafter. On day 14 after engraftment HkiL cell progeny with a GC phenotype were present in both DOC- and DOC-LAV-infected 10-cell-recipients, offering an opportunity for a comparative analysis of transcriptional signatures associated with attrition. We purified HkiL GC B cells from DOC- and DOC-LAV-viremic recipients by FACS sorting on day 14 (Fig. S4A,B) and processed them for bulk RNAseq. A gene set enrichment analysis on the MSigDB hallmark pathways revealed that target genes of c-Myc, which is downregulated in GC dark zone cells ^104^, were expressed at lower levels when HkiL GC B cells responded to DOC infection as compared to DOC-LAV infection (Fig. 4F). Consistent therewith, the comparison to a published gene set for GC dark zone vs. light zone differentiation^104^ indicated that HkiL GC B cells in DOC-infected animals exhibited a more pronounced dark zone signature than their counterpart in DOC-LAV-infected mice (Fig. 4G,H, S4C). This dark zone bias of Bcl6^+^B220^+^ HkiL GC B cells from DOC-infected animals was also reflected in an increased proportion of CXCR4^+^CD86^-^ dark zone (DZ) cells and a reduced fraction of CXCR4^-^CD86^+^ light zone (LZ) cells in flow cytometry (Fig. 4I-K, S4D). Efficient GC reactions require B cells to attenuate their BCR signaling and the failure to do so can result in a dark zone bias ^105,106^. One pathway to attenuated BCR signaling comprises the remodeling of the surface glycan composition on GC B cells ^105^, which can be detected by means of the widely used GL7 antibody. Interestingly, Bcl6^+^B220^+^ HkiL GC B cells from DOC-infected recipients exhibited lower levels of GL7 than those from DOC-LAV-infected controls (Fig. 4L). Taken together, these findings suggested that persistent viral infection resulted in phenotypic alterations that preferentially affect high-affinity GC B cells.

**Figure 4:**
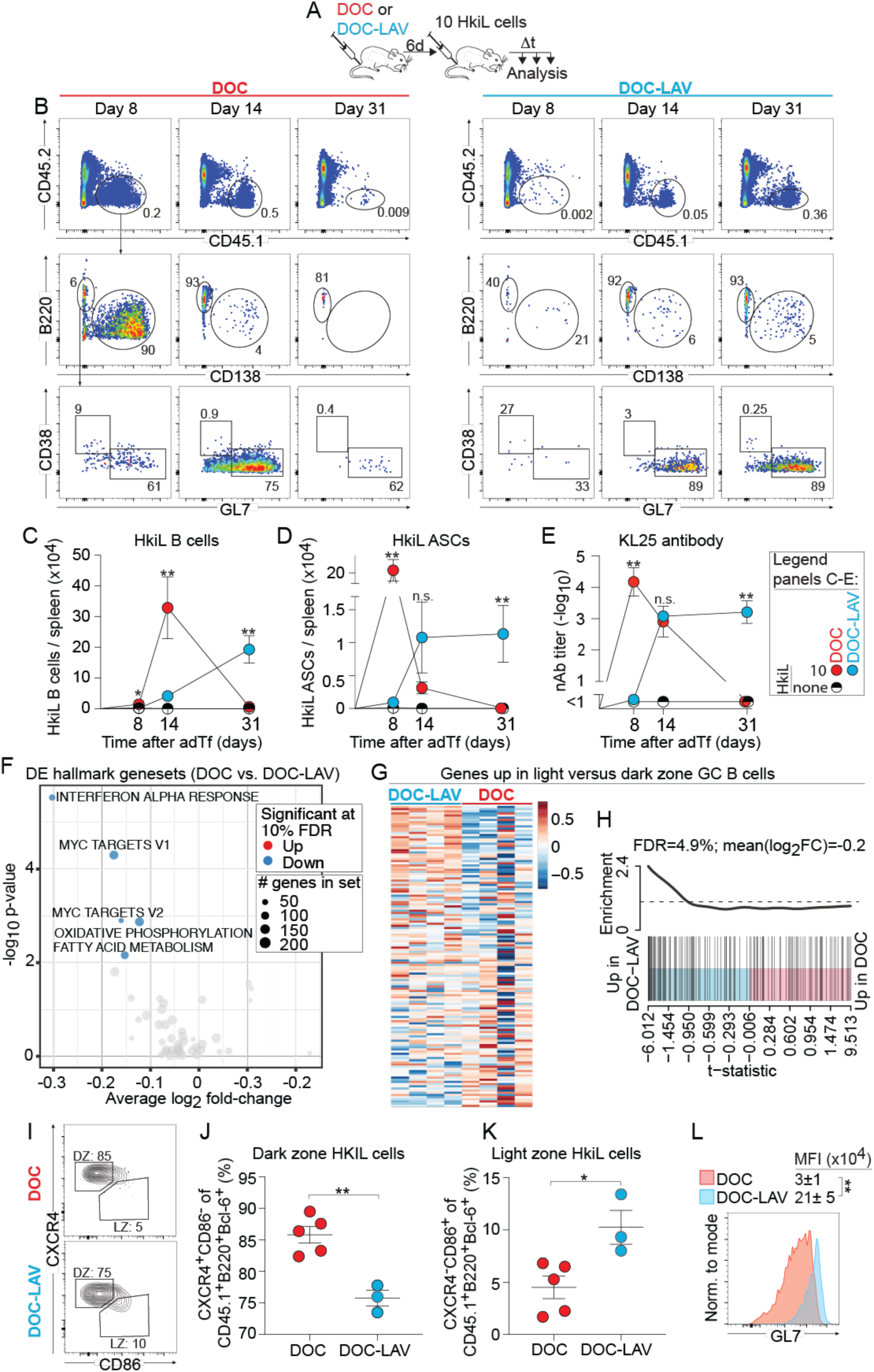
On their path to attrition high-affinity B cells pass through an ASC burst and a subsequent dark zone-biased GC stage. A-E: We infected mice with DOC or DOC-LAV on d-6 and on d0 engrafted either 10 HkiL cells or none (”none”, noTF) (A). HkiL progeny in spleen were analyzed on d8, d14 and d31. Representative FACS plots (B; n(d8, d14, d31)=4, 4 and 3 mice for DOC; n(d8, d14, d31)=3, 3 and 5 mice for DOC-LAV; n(d8, d14, d31)=1, 1 and 3 mice for noTf), pre-gated on live lymphocytes (Fig. S1A). Splenic HkiL progeny (CD45.1^+^, B: top) were analyzed for expression of B220 and CD138 (B: center), and their B220^+^CD138^—^ subset was analyzed for GL7 and CD38 expression (B: bottom). CD45.1^+^B220^+^CD138^-^ HkiL progeny of B cell phenotype and CD45.1^+^B220^-^CD138^+^ HkiL progeny of ASC phenotype were enumerated (C,D), and (HkiL cell-derived) nAb titers in serum (E) were determined over time. F-G: In an experiment using GFP-transgenic HkiL cells and conducted as in (A) we sorted on d14 live CD45.1^+^B220^+^CD138^-^GFP^+^ HkiL B cell progeny (Fig. S4A). Total RNA of the sorted cells was processed for bulk RNAseq (F-H). Differentially expressed (DE) hallmark gene sets with an FDR<0.1 are shown in (F). Heat maps display the expression of genes reported to be upregulated in light zone (LZ) as compared to dark zone (DZ) GC B cells^104^ (see also Fig. S4C). Each column of the heatmap represents an individual mouse (n=4 per group). Pair-wise self-contained gene set testing (H). I-L: In an experiment conducted as in (A) we analyzed HkiL progeny in spleen on d17. Representative FACS plots (n=5 for DOC, n=3 for DOC-LAV), pre-gated on live CD45.1^+^B220^+^Bcl6^+^ HkiL cells (Fig. S4D) showing their LZ (CD86^+^CXCR4^-^) and DZ (CD86^-^CXCR4^+^) distribution (I), proportional repartition into these zones (J,K) and GL7 expression profile (L). Representative histogram plots pre-gated on live CD45.1^+^CD19^+^Bcl6^+^ HkiL GC B cells (Fig. S4D). Numbers in FACS plots indicate the percentage of gated cells as the group mean (B, I). Symbols and error bars in (C,D,E) show the mean±SEM. For statistical analysis, cell numbers in (C,D) were log-converted and samples without any detectable HkiL cells were assigned the highest count recorded in the noTf group. For each time point the values of DOC- and DOC-LAV-infected mice were compared by unpaired Student’s t-tests with Bonferroni correction. Two-way ANOVA with Bonferroni’s multiple comparison test was performed in (E) and for each time point significant differences between DOC- and DOC-LAV-infected mice are reported. Unpaired Student’s t tests were performed in J-L. Data in (B-E, I-L) show one representative out of two similar experiments.*: p<0.05; **: p<0.01; n.s.: not statistically significant.

### Attrition of high-affinity B cells in chronic viral infection is Blimp-1-dependent

In addition to the above transcriptional alterations HkiL GC B cells from DOC-infected mice exhibited a more pronounced Blimp-1 signature than those responding to DOC-LAV. More specifically, genes known to be activated by the transcription factor Blimp-1 ^107^ (encoded by the *Prdm1* gene) were higher in DOC- than in DOC-LAV-infected mice, whereas Blimp-1-repressed genes exhibited the opposite pattern (Fig. 5A-D). Blimp-1 is a master transcription factor of plasma cell differentiation and can also play an important role in the transcriptional regulation of GC B cells ^108-112^, prompting us to test whether Blimp-1 expression in B cells was required for attrition. Blimp-1 deficiency is embryonically lethal, but fetal liver cells from animals homozygous for a functional null allele of Prdm1 (encoding for Blimp-1; *Prdm1*^-/-^ mice) can be used to reconstitute irradiated recipients and give rise to mature B cells ^108^. Following this strategy we generated donors of either Blimp-1-deficient or -sufficient HkiL cells (HkiL-*Prdm1*^-/-^, HkiL-*Prdm1*^wt/wt^) from which we transferred 10 cells into DOC-infected recipients. When analyzed 14 days later, progeny of both types of cells were readily detectable (Fig. 5E-G). As expected, HkiL-*Prdm1*^-/-^ progeny were virtually devoid of CD138^+^B220^low/int^ ASCs, they failed to produce detectable amounts of KL25 serum antibody and by consequence the corresponding recipients had higher viral loads than those engrafted with HkiL-*Prdm1*^wt/wt^ cells (Fig. 5F,H,I). When recipients were analyzed 24 days after engraftment, HkiL-*Prdm1*^wt/wt^ cells had undergone attrition, as expected (Fig. 5F,G). In contrast, HkiL-*Prdm1*^-/-^progeny persisted in DOC-infected recipients in numbers comparable to those determined on day 14, thus offering no evidence of attrition. Unlike in the context of chronic DOC infection, Blimp-1-deficient and -sufficient HkiL cells expanded and persisted in similar numbers when triggered by rPICV vaccination (Fig. S5C-F). Hence, Blimp-1 deficiency prevented the attrition of high-affinity B cells in chronic viral infection but did not afford a clear advantage to B cells responding to vaccination. Taken together, the resistance of Blimp-1-deficient B cells to attrition suggested the latter was the consequence of a cell-intrinsic reprogramming of GC B cells in the context of chronic viral infection.

**Figure 5:**
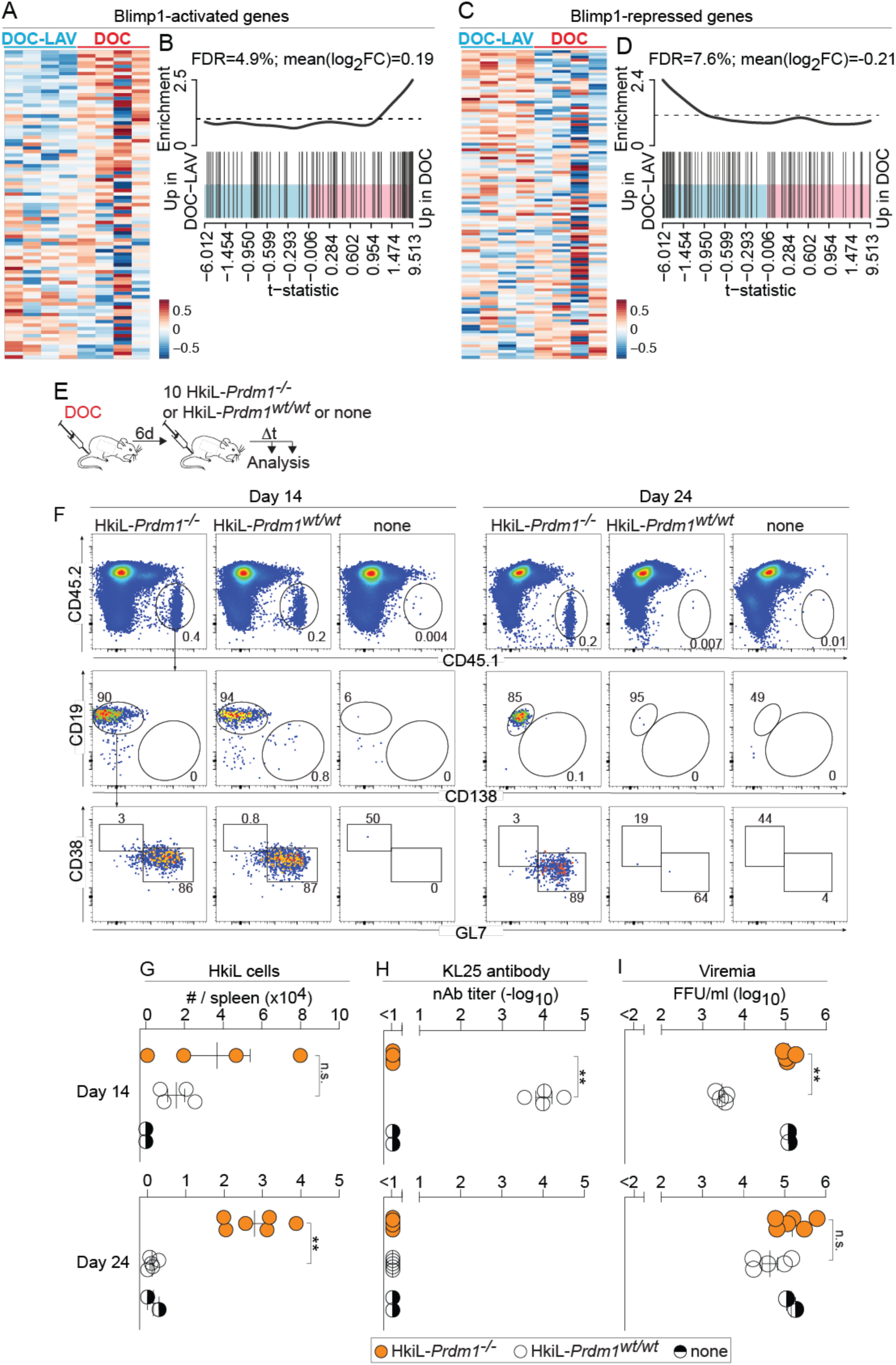
Attrition of high-affinity B cells in chronic viral infection is Blimp-1-dependent. A-D: The data sets from Fig. 4F-H were analyzed for Blimp-1-activated (A,B) and Blimp-1-repressed genes (C,D)^107^. See also Fig. S5A,B. Each column of the heatmaps in (A,C) represents an individual mouse (n=4 per group). Pair-wise self-contained gene set testing is shown in (B,D). E-I: We infected mice with DOC on d-6, on d0 we engrafted either 10 HkiL-*Prdm1*^-/-^ or 10 HkiL-*Prdm1*^wt/wt^ cells or none (noTF) and we analyzed these cells’ progeny in spleen on d14 and on d24. Representative FACS plots (F; d14: 4 HkiL-*Prdm1*^-/-^ recipients, 4 HkiL-*Prdm1*^wt/wt^ recipients, 2 noTf; d24: 6 HkiL-*Prdm1*^-/-^ recipients, 5 HkiL-*Prdm1*^wt/wt^ recipients, 2 noTf), pre-gated on live lymphocytes (Fig. S1A). HkiL progeny (CD45.1^+^, top) were analyzed for expression of CD19 and CD138 (center), and their CD19^+^CD138^—^ subset was analyzed for GL7 and CD38 expression (bottom). HkiL cell progeny (G), KL25 serum antibody (H) and viremia (I) were determined on d14 and d24. Numbers in FACS plots indicate the percentage of gated cells as the group mean (F). Values of HkiL-*Prdm1*^-/-^ and HkiL-*Prdm1*^wt/wt^ recipients in G-I were compared by unpaired Student’s t tests. Results in (E-I) show one representative out of two experiments. **: p<0.01; n.s.: not statistically significant.

## Discussion

The present findings in the context of chronic LCMV infection indicate that antiviral B cells with a high-affinity receptor mount an abortive response and are eliminated by attrition unless they can get the upper hand over the virus by either clearing it or forcing it into mutational escape. The naïve precursors of LCMV- and HIV-nAb-producing B cells in mice and humans, respectively, are, however, mostly of low to intermediate affinity, and substantial hypermutation is required for these cells to reach affinity ranges that confer neutralizing capacity^32,53-55,101^. Our observations predict on the one hand that the average nAb-producing B cell precursor will efficiently expand and affinity-mature in chronically infected individuals. On the other hand our data suggest that each clone’s highest-affinity progeny will be pruned by attrition, thus continuously ridding the antiviral B cell response of its most proficient output.

Exogenously supplied as well as endogenously produced antibodies can modulate antigen- and/or epitope-specific GC B cell responses by so-called “antibody feedback” i.e. by blocking B cell access to antigen ^98-100^. While commonly known for its negative regulatory effects in the context of vaccination i.e. under conditions of limited antigen availability, the present findings raise the possibility that attenuation of BCR signaling by antibody feedback may benefit GC B cell responses under conditions of chronic high-level viremia. Accordingly, the protective effect of AAV-WEN3 therapy against HkiL cell attrition in DOC infection may help explain the observation that bnAb treatment of HIV-viremic individuals augmented their endogenous HIV-nAb response^113^ and, analogously, that LCMV-nAb therapy improved polyclonal virus-specific GC B cells responses in chronically infected mice^90^. The molecular mechanism whereby AAV-WEN3 therapy attenuates HkiL cell attrition requires, however, further investigation. Besides attenuation of BCR signaling as discussed above, the viral load reduction afforded by the passively administered nAb may exert additional beneficial effects on HkiL cells e.g. by reducing detrimental consequences of virus-induced inflammation ^29-31,97^. Also, the mechanism whereby Blimp-1 deficiency protects HkIl cell from attrition warrants further investigation. Blimp-1 negatively regulates cell-cycle progression and c-Myc expression by GC B cells ^112,114^. In light of repressed c-Myc transcriptional signatures in HkiL cells destined to attrition it seems plausible that Blimp1 deficiency counteracted attrition by enabling the cells to maintain critical c-Myc levels and to more efficiently progress in cell cycle. Not mutually exclusively, Blimp1-mediated downregulation of Bcl6 and upregulation of IRF4 ^112^ may have promoted attrition as suggested by the massive early ASC burst typical of the HkiL cell response to DOC.

Of note in this context, ASC-biased B cell compartments represent a long-standing observation in HIV-viremic individuals and have been associated with subversion of humoral immunity^115,116^. While preferential ASC differentiation of high-affinity B cells serves presumably to ascertain the effectiveness of the serum antibody responses in the context of acute infection and vaccination^110,117^ we propose that in chronic viral infection the same may culminate in attrition, thus promoting humoral immune subversion.

Our study with its experimental set-up has clear limitations. LCMV infection of mice represents an imperfect surrogate of human persistent viral diseases despite having contributed concepts of broad utility such as the exhaustion of CD8 T cells in persistent viral infection, their reinvigoration by PD-1-targeted therapy and mutational escape from CD8 T cell control^8-17,118^. Moreover, our study relied on the adoptive transfer of a monoclonal population of receptor knock-in B cells. While representing the only experimental approach whereby to control key parameters such as the affinity of antiviral B cells and the timing of their recruitment into the immune response, the possibility remains that other B cell clones in other infection settings would respond differently. Observations with adoptively transferred HBV-specific B cells in a mouse model of persistent HBV infection indicate, however, that a plasmablast burst followed by the disappearance of virus-specific B cells from the GC is not unique to HkiL cells in chronic LCMV infection^119^.

Taken together, this study describes the affinity-dependent clonal attrition of specific B cells during viral persistence, suggesting the chronic infection context perturbs affinity hierarchies of antiviral B cell responses. The understanding of humoral immune subversion by persisting viruses with molecular insights into the underlying mechanisms may open new avenues for therapeutic interventions aimed at improving humoral immune control of chronic microbial diseases.

## Acknowledgments

We wish to thank Katrin Martin and the entire Experimental Virology lab for helpful discussions, Cynthia Saadi for excellent technical assistance with immunohistochemistry, the entire Genomics Facility Basel for RNAseq, Cécile Cumin and the entire DBM flow cytometry core facility for FACS-sorting, Stephen Nutt for generously providing *Prdm1*^-/wt^ mice. Bioinformatic analysis calculations were performed at sciCORE (http://scicore.unibas.ch/) scientific computing center at University of Basel.

## Author contributions

M.D, T.A.M., J.F., W.V.B., A.F.M., A.L.K., Y.I.E., M.F., M.C., K.N., K.M., J.R., Ph.D., D.M. and D.D.P. designed experiments. M.D., T.A.M., J.F., W.V.B., A.F.M., M.L., K.T., A.L.K., Y.I.E, M.F., M.C., K.S., J.R., Ph.D., I.W. performed experiments. M.D., T.A.M., J.F., W.V.B., A.F.M., A.L.K., Y.I.E., J.R., Ph.D., I.W. and D.D.P. analyzed data. M.D. and D.D.P. wrote the manuscript.

## Methods

### Data availability

- scRNAseq data are deposited at GEO under GSE296780.
- Raw data of the experimental results reported in this study will be deposited with Zenodo and will be publicly available as of the date of publication under DOI 10.5281/zenodo.15600975.

### Mice and animal experiments

C57BL/6J wildtype mice were originally purchased from Charles River and were bred locally for experiments. HkiL mice carrying both a B cell receptor (BCR) heavy and light chain V(D)J knock-in originating from the LCMV-neutralizing GP-1-specific monoclonal antibody (mAb) KL25 have been described ^32^. TgL mice were used as recipients for HkiL B cells to avoid anti-idiotypic rejection^29,32,101,120^. B cell-deficient JHT mice^121^ and interferon receptor deficient mice (*ifnar*^-/-^)^122^ were obtained from the Swiss Immunological Mouse Repository (SwImMR). HkiL mice were crossed to *ifnar*^-/-^ mice to obtain HkiL-*ifnar*^-/-^ cell donors (CD45.1^+^). As recipients of HkiL-*ifnar*^-/-^ and control HkiL B cells in the respective experiments we used *ifnar*^-/-^ mice that were intercrossed with TgL mice (for simplicity referred to as *ifnar*^-/-^ mice in the manuscript’s text). HkiL mice were intercrossed with *Prdm1*^-/wt^ mice^108^ with a GFP knock-in disrupting the *Prdm1* locus, and separately with UBI-GFP mice (C57BL/6-Tg(UBC-GFP)30Scha/J; Jax #004353) to obtain HkiL-*Prdm1*^gfp/wt^ mice and HkiL-GFP mice, respectively (both CD45.1^+^). HkiL, HkiL-GFP and HkiL-*Prdm1*^gfp/wt^ mice were kept on a RAG2-deficient background^32^. To prevent rejection of GFP-expressing HkiL-*Prdm1*^-/-^ B cells (see below) and control HkiL cells in the respective experiments we used TgL mice with liver-specific expression of EYFP. For this, TgL mice were intercrossed with Albumin-Cre (B6.Cg-Speer6-ps1Tg(Alb-cre)21Mgn/J; Jax #003574) and R26-stop-EYFP mice (B6.129X1-Gt(ROSA)26Sortm1(EYFP)Cos/J; Jax #006148). All mouse lines were on a C57BL/6J background and were bred at the ETH Phenomics Center (EPIC) of the Swiss Federal Institute of Technology Zurich or at the University of Basel mouse breeding facility under specific pathogen-free (SPF) conditions. Experiments were performed at the University of Basel in accordance with the Swiss Law for animal protection and with authorization from the Veterinary Office Basel Stadt. Adult animals from both genders were used to reduce the number of animals bred for research purposes. Within each experiment the animals were age- and sex-matched. Experiments were not conducted in a blinded fashion.

### Generation of HkiL-*Prdm1^-^*^/-^ and HkiL-*Prdm1*^wt/wt^ control B cell donors

HkiL-*Prdm1*^-/wt^ mice homozygous for the B cell receptor (BCR) heavy and light chain V(D)J knock-ins were mated with *Prdm1*^-/wt^ mice and 48 hours later the males were removed from the breeding cages. Fourteen days after mating the females were sacrifices, embryos were removed, their livers were extracted, placed in DPBS supplemented with 3% FCS and kept on ice. A piece of each embryo’s front paw was processed for DNA extraction using the QIAmp Fast DNA tissue kit (Quiagen). Upon identification of *Prdm1*^-/-^ and *Prdm1*^wt/wt^ control embryos by TaqMan PCR (see below), the respective liver samples were filtered through a 70 μm cells strainer and combined with JHT bone marrow for intravenous implantation into lethally irradiated (11 Gy one day beforehand) C57BL/6J recipients. Reconstitution of the recipients with HkiL-*Prdm1*^-/-^ or HkiL-*Prdm1*^wt/wt^ control B cells (CD45.1^+^) of fetal liver cell origin was verified in the respective recipient’s blood ≥30 days later, and successfully reconstituted mice were used as B cell donors for adoptive transfer experiments in Fig. 5E-I. For the experiment reported in Fig. S5C-F splenic B cells from a non-chimeric HkiL mouse were used as HkiL- *Prdm1*^wt/wt^ control cells.

### TaqMan PCR-based genotyping of HkiL-*Prdm1^-/-^* and HkiL-*Prdm1*^wt/wt^ embryos

To identify HkiL-*Prdm1*^-/-^ and HkiL-*Prdm1*^wt/wt^ control embryos and to differentiate them from HkiL-*Prdm1*^-/wt^ animal we performed TaqMan PCR analyses on DNA extracted from embryonic biopsies as described above. In brief, the extracted DNA was quantified by optical density measurements and the concentration of samples was adjusted to 50 ng/μl. The samples were then analyzed in duplicates on a StepOnePlus Real-Time PCR System 96-well device (Applied Biosystems) using the following primer-probe sets. The unmodified *Prdm1* locus was amplified using primers 5’-CTTGGCTGCCAGGCAGTGC-3’ and 5’-GCCAGGTATGTACAATGCAGATGC-3’ with a probe 5’-FAM-TTGAGCCATAGGAGACC-BHQ1-3’. The *Prdm1*-ko locus was detected by amplification of the inserted GFP sequence using primers 5’-CTCGTGACCACCTTGACCTA-3’ and 5’-GAAGTCGTGCTGCTTCATGT-3’ with the probe 5’-FAM-CGGACGAAGCACTGCACGCCG-BHQ1-3’. To normalize for DNA content in the reactions we used ApoB-binding primers 5’-CACGTGGGCTCCAGCATT-3’ and 5’-TCACCAGTCATTTCTGCCTTTG -3’ with the probe 5’-FAM-CCAATGGTCGGGCACTGCTCAA-BHQ1-3’.

### Cells, viruses, virus titration and infection of mice

NIH 3T3 cells (CRL-1658) and MDCK cells (CCL34) were purchased from ATCC. BHK21 cells (85011433) and VeroE6 cells were both from ECACC (# 85020206). All cells were cultured at 37 °C in an atmosphere of 5 % CO_2_. They were regularly tested for mycoplasma and confirmed negative. The genetically engineered LCMV clone 13-based rCl13/WE and the Armstrong-based rARM/WE have been described^84,123^. rARM/WE-LAV was generated from cDNA as previously described^124^. The wildtype LCMV strain Docile^93^ (DOC) was originally obtained from Rolf Zinkernagel. A cDNA rescue system for DOC was built using as a basis a published set of plasmids^92^ that were generously provided by Dimitrios Moskophidis. Modifications to the published viral sequence^92^ were made as outlined in Fig. S2B. cDNA-derived DOC of wildtype sequence as well as DOC with the N119S point mutation in its GP (DOC-LAV) were generated following established procedures^124^ and were used throughout the study except for the wildtype DOC isolate used in the experiment in Fig. S2B as indicated. Pichinde- (PICV-) based viruses expressing instead of their natural envelope glycoprotein the LCMV strain WE glycoprotein (rPICV) or its N119S-mutant (rPICV-LAV) were engineered as described^96^. Clone 13- and Armstrong-based viruses were propagated on BHK-21 cells, DOC-based viruses on MDCK cells. For infection of mice rCl13/WE, DOC and DOC-LAV were administered intravenously at a dose of ≥2x10e6 focus-forming units (FFU), rARM/WE and rARM/WE-LAV were administered intraperitoneally at a dose of 200 FFU. rPICV and rPICV-LAV were administered intravenously at a dose of 10e6 FFU. For the determination of viremia one drop of blood was collected into 950 μl of BSS supplemented with 1 IE/ml heparin (Na-Heparin, Brown). Infectious titers in viral stocks and blood samples of animals infected with LCM viruses or immunized with PICV-based viruses were determined by immunofocus assays on NIH 3T3 cells as described^96,125^.

### Adeno-associated viral vectors and their administration to mice

An adeno-associated viral (AAV) vector genome expressing the LCMV-neutralizing mAb WEN3^81,101,126^ (AAV-WEN3) was designed and generated following established strategies and procedures^90,127^. AAV vector particles with an AAV8 capsid were produced and titrated by the Viral Vector Facility (VVF) of the University of Zurich, Switzerland. AAV-WEN3 was administered to mice at a dose of 10e11 vector particles (v.p.) in a total volume of 40 µl into both thigh muscles.

### Adoptive B cell transfer

For adoptive transfer of B cells from HkiL and HkiL-GFP mice as well as from HkiL-*Prdm1*^-/-^ and HkiL-*Prdm1*^wt/wt^ chimeras, splenic single cell suspensions were generated by mechanical disruption and B cells were purified using the EasySep mouse B cell isolation kit (STEMCELL, #19854) following the manufacturer’s instructions. The cells were adoptively transferred into the tail vein in either BSS or RPMI. Assuming a splenic take of ∼5% (^80,82,83^; Fig. S1), we administered 200 purified B cells to engraft 10 cells into the spleen of recipients (10-cell-recipients), 600 B cells to generate 30-cell recipients, 2,000 B cells for 100-cell recipients and 20,000 cells for 1000-cell-recipients. The group of mice labeled as “100-cell” recipients in Fig. 1K-O summarizes two groups of mice given 2000 and 5000 cells, respectively, to engraft 100 or 250 HkiL cells.

### Flow cytometry

To obtain single cell suspension, spleens were mechanically disrupted using a metal grid and syringe plunger. The cells were then resuspended in RPMI (Sigma, R2405) supplemented with 20mM HEPES, 5mM MgCl_2_, 1mM NaPyruvate, MEM non-essential amino acids, 5% FCS, Penicillin / Streptomycin and 0.05 mM β-mercaptoethanol. Media and buffers were adjusted to mouse osmolarity using NaCl^128^. Lymph nodes were mechanically disrupted using a syringe plunger and a 70 μm cells strainer and resuspended in FACS buffer (PBS supplemented with 10% Opti-MEM, 1% FCS, penicillin / streptomycin, 10 mM HEPES, 5 mM NaCl, 1 mM Na-Pyruvate, MEM non-essential amino acids, 0.05 mM β-mercaptoethanol and 1 g/l D-Glucose). Bone marrow was flushed out of tibiae using a syringe and then was processed as for lymph node samples. Prior to staining, spleen and lymph nodes suspensions were incubated for 10 min at room temperature (RT) with DNAse I (50 μg/ml; Roche, 10104159001) in FACS buffer.

Stains were performed in FACS buffer supplemented with polyclonal rat IgG and anti-mouse CD16/CD32 (2.42; BioXcell, BE0307) to block nonspecific antibody binding. When more than one brilliant violet dye was in the staining mix, a BD HorizonTM brilliant stain buffer (BD Biosciences, 566385) was added to the staining mix. Antibodies against CD45.1 (A20), CD45.2 (104), CD45R/B220 (RA3-6B2), CD138 (281-2/2PH1), CD19 (6D5 or 1D3), Bcl-6 (K112-91), CD38 (REA616 or 90), GL7 (GL7), CXCR4 (2B11 or L276F12), CD86 (GL1 or PO3), Ter119 (ter119), CD95 (JO2), the Zombie U.V. Fixable Viability Kit and the Zombie NIR Fixable Viability Kit were purchased from BD Biosciences, Miltenyi Biotec, Biolegend, and Thermofisher (Invitrogen, ebiosciences) and used for staining. Samples were acquired on BD LSRFortessa (BD Biosciences) and Aurora (Cytek) flow cytometers and were analyzed using FlowJo (BD Biosciences).

### Determination of GP1-binding and LCMV-neutralizing antibody concentrations in mouse serum

For serum collection we used Multivette 600 Serum gel tubes (Sarstedt, 15.1674) following the manufacturer’s instructions. For enzyme-linked immunosorbent assays (ELISA) we followed established procedures^102^. In brief, 96-well flat-bottom high-binding well plates were coated overnight at 4°C with anti-human-Fc antibody diluted in coating buffer (15mM Na_2_CO_3_, 35mM NaHCO_3_ in ddH_2_O, pH 9.6). The next day, the plates were blocked with 5% milk PBS-0.05% Tween-20 (PBST) for 1h at RT. The blocking buffer was flicked off and GP1-Fc fusion protein was added to the plates for 1h at RT followed by three washes with PBST. In a separate 96-U-bottom plate we prepared 3-fold serial dilutions of serum samples in PBST supplemented with 1% FCS, then we transferred the diluted serum into the prepared 96-well flat-bottom high-binding well plates, followed by a 1h incubation at RT. Plates were washed 3 times with PBST and incubated with secondary HRP-couples antibody in PBS supplemented with 1% FCS for 1h at RT. Plates were then washed 3 times with PBST and once with PBS and an ABTS color reaction mix was added consisting of 0.5 mg/ml ABTS (Thermo Fischer, 34026), 28 mM Na_2_HPO_4_ and 0.1 % H_2_O_2_ in H_2_O and developed for 30 min. We stopped the color reaction by adding 1% sodium dodecyl sulfate (SDS) in ddH2O and measured optical density (O.D.) at a wave length of 405 nm using an Infinite M Plex reader (Tecan). To calculate serum concentrations of GP1-specific KL25 antibody produced by adoptively transferred HkiL cells we used KL25 monoclonal antibody standard. Naïve control serum samples were used as negative controls.

To determine neutralizing antibody titers in serum, samples from infected mice and naïve negative control sera were pre-diluted ten-fold and then three-fold serial dilutions were prepared in 96-well flat-bottom tissue culture plates using MEM supplemented with 2% FCS as diluent. Monoclonal KL25 antibody was included as a positive control. The sample-containing plates were irradiated in a UV chamber for 5 min to inactivate residual infectivity in mouse serum. Then the diluted serum samples were incubated with an equal volume of medium containing approximately 1000 infectious units of green fluorescent protein-expressing replication-deficient vesicular stomatitis virus (rVSV-EGFP^129^, generously provided by Gert Zimmer) pseudotyped with LCMV-WE glycoprotein for 1.5h at 37°C. Next, the plates were incubated with Vero E6 cells (3×10^4^ cells/well) for 24h at 37°C and fixed with 1% PFA. After fixation, PFA was flicked off and PBS was added until quantifying green cells using an Immunospot S6 device (C.T.L.). The 50% neutralization titer (NT_50_) was calculated using 4-parameter non-linear regression to determine the serum dilution yielding a 50% reduction of viral infectivity. Inverted titers were log-converted and are displayed in logarithmic format.

### Viral sequence determination

For viral RNA isolation and sequencing, BHK21 cells were infected with viremic mouse blood samples and supernatant was collected two days later. Viral RNA was extracted from virions using the QIAamp Viral RNA Mini Kit (QIAGEN) in accordance with manufacturer’s instructions. cDNA synthesis and PCR amplifications of the DOC GP region was performed with the OneStep RT-PCR kit (QIAGEN) according to the manufacturer’s protocol and using the primers 5’-ATGGGCCAAATTGTGACAATGTT-3’ and 5’-CAGCGTCTTTTCCAGATAGTTT-3’. Alternatively we performed reverse transcription with the SuperScript IV first strand synthesis system (Thermofisher), followed by PCR using Phusion polymerase (New England Biolabs). Primers 5’-CGCACAGTGGATCCTAGGC-3’ and 5’-GGGTGAGTTAGCTACAGGTTTC-3’ were used to amplify the DOC wildtype NP sequence. The PCR products were run on 1.5% agarose gel, amplicons of the correct size were excised, purified using the QIAquick Gel Extraction Kit (QIAGEN) and subject to Sanger sequencing (Microsynth AG, Switzerland).

### FACS-sorting and bulk RNA sequencing of HkiL cells

For bulk RNA sequencing, HkiL-GFP cells from DOC- and DOC-LAV-infected mice were bead-enriched using anti-CD45.1-PE antibody and the EasySep^TM^ PE Positive Selection Kit II (STEMCELL, 17684) according to the manufacturer’s instructions. Subsequently HkiL B cells (CD45.1^+^GFP^+^B220^+^CD138^-^ cells) were sorted directly into lysis buffer (TAKARA) using a FACSAriaII (Beckton Dickinson) and stored at -80°C until further use. Library preparation was performed from the direct lysis of 300 to 500 cells in 10x Lysis Buffer (Cat# 635013, Takara Bio) and Recombinant RNase Inhibitor (Cat# 2313A, Takara Bio) as described in the kit SMART-Seq Stranded Kit (Cat# 634444, Takara Bio) in conjunction with the SMARTer RNA Unique Dual Index Kit (Cat# 634452, Takara Bio) to produce ribosomal depleted libraries. The same conditions were applied for all samples with a standard time of 6 min for fragmentation, 5 cycles for PCR 1 and 14 for PCR 2, and a single final cleanup cycle. Libraries were quality-checked on the Fragment Analyzer (Advanced Analytical, Ames, IA, USA) using the High Sensitivity NGS Fragment Analysis Kit (Cat# DNF-474, Advanced Analytical) revealing excellent quality of libraries (average concentration was 16±10 nmol/L and average library size was 408±26 base pairs). Samples were pooled to equal molarity. The pool was quantified by Fluorometry using the QuantiFluor ONE dsDNA System (Cat# E4871, Promega, Madison, WI, USA) and sequenced on an Illumina NextSeq 500 instrument using the NextSeq 500 High Output Kit 75-cycles (Illumina, Cat# FC-404-1005) to produce paired-end 38nt reads (in addition: 8 bases for index 1 and 8 bases for index 2). Flow lanes were loaded at 1.8pM of pool with 1% PhiX. This Nextseq runs compiled a large number of reads (on average per sample: 42±4 millions pass-filter reads).

### Analysis of bulk RNA sequencing data

Data analysis was performed by the Bioinformatics Core Facility, Department of Biomedicine, University of Basel. Read quality was assessed with the FastQC tool (version 0.11.9). Reads were mapped to the mouse genome mm39 with STAR^130^ (version 2.7.10a) with default parameters, except filtering out multimapping reads with more than 10 alignment locations (outFilterMultimapNmax=10) and filtering reads without evidence in the spliced junction table (outFilterType=“BySJout”). The *featureCounts^131^* function from the Rsubread package (version 2.0.6) was used to count the number of reads (5’ ends) overlapping with the exons of each gene (Ensembl release 110 genes) assuming an exon union model. All subsequent analyses were performed using the R software (version 4.3.1) and Bioconductor^132^ 3.18 packages. A total of 19,417 genes with CPM values above 1 in at least 3 samples (n-1 the number of biological replicates), and with a protein-coding gene biotype, were retained for the differential expression analysis. To account for complexity differences commonly observed in RNAseq libraries from low-input material, the function voomWithQualityWeights^133,134^ from the *limma* package (version 3.58.1) was used (with a cyclic loess normalization ^135^) to combine observational-level with sample-specific quality weights prior to differential expression analysis comparing DOC and DOC-LAV conditions. P-values were adjusted by controlling the false discovery rate (FDR; Benjamini-Hochberg method) and genes with a FDR lower than 5% were considered significant. Gene set enrichment analysis was performed with the function *camera^136^* from the limma package (using the default parameter value of 0.01 for the correlations of genes within gene sets) using gene sets from the mouse collections of the MSigDB Molecular Signatures Database^137^ (version 2023.2), notably the hallmark gene sets collection. A collection of custom gene sets was also collected from previously published studies and datasets. We filtered out sets containing less than 10 genes, and gene sets with a FDR lower than 10% were considered significant.

### Immunohistochemistry and Image analysis

For fluorescent immunohistochemistry spleen samples were fixed with HEPES-glutamic acid buffer-mediated organic solvent protection effect (HOPE, DCS Innovative, HL001R2500), subsequently embedded into paraffin and cut into 2 μm thick sections using a microtome. The CD45.1 congenic marker of HkiL cells was stained using the primary anti-mouse CD45.1-FITC antibody (A20, Invitrogen #11-0453-85) and secondary rabbit anti-FITC antibody (Invitrogen #71-1900), followed by tertiary Alexa Fluor 488 donkey anti-rabbit antibody (Life Technologies #A21206). Naïve B cells were visualized using rat anti-mouse IgD monoclonal antibody (Bioegend, clone 11-26c-2a, #405702), which had been labelled in-house with Alexa Fluor 647 (Invitrogen Labeling kit #A20186). GCs were stained with AF568-labelled peanut agglutinin (Life Technologies #L32458) and nuclear staining was performed using 4ʹ,6-diamidino-2-phenylindole (DAPI, Invitrogen). For image acquisition, slides were scanned using a Pannoramic 250 FLASH II (3DHISTECH) Digital Slide Scanner at 20x magnification.

### Statistical analysis

For statistical analysis we used GraphPad Prism Software (Version 10.2.3). For the comparison of a single parameter between two groups we performed unpaired two-tailed Student’s t-tests. For the comparisons of one parameter between multiple groups we performed 1-way ANOVA with Bonferroni’s or Tukey’s post-test. For the comparison of multiple groups against a reference control group one-way ANOVA with Dunnett’s post-test was performed. Repeated measures ANOVA was performed for a pairwise comparison of time courses. To analyze repeated measurements of two or more groups we used two-way ANOVA with Tukey’s or Bonferroni’s post-test. p≤0.05 was considered not statistically significant (n.s.), p<0.05 as statistically significant (*) and p<0.01 as highly significant (**). Absolute cell counts, where indicated in legends, as well as antibody titers and viral loads were log-converted to obtain a near-normal distribution for statistical analysis.

---

**Figure S1:**
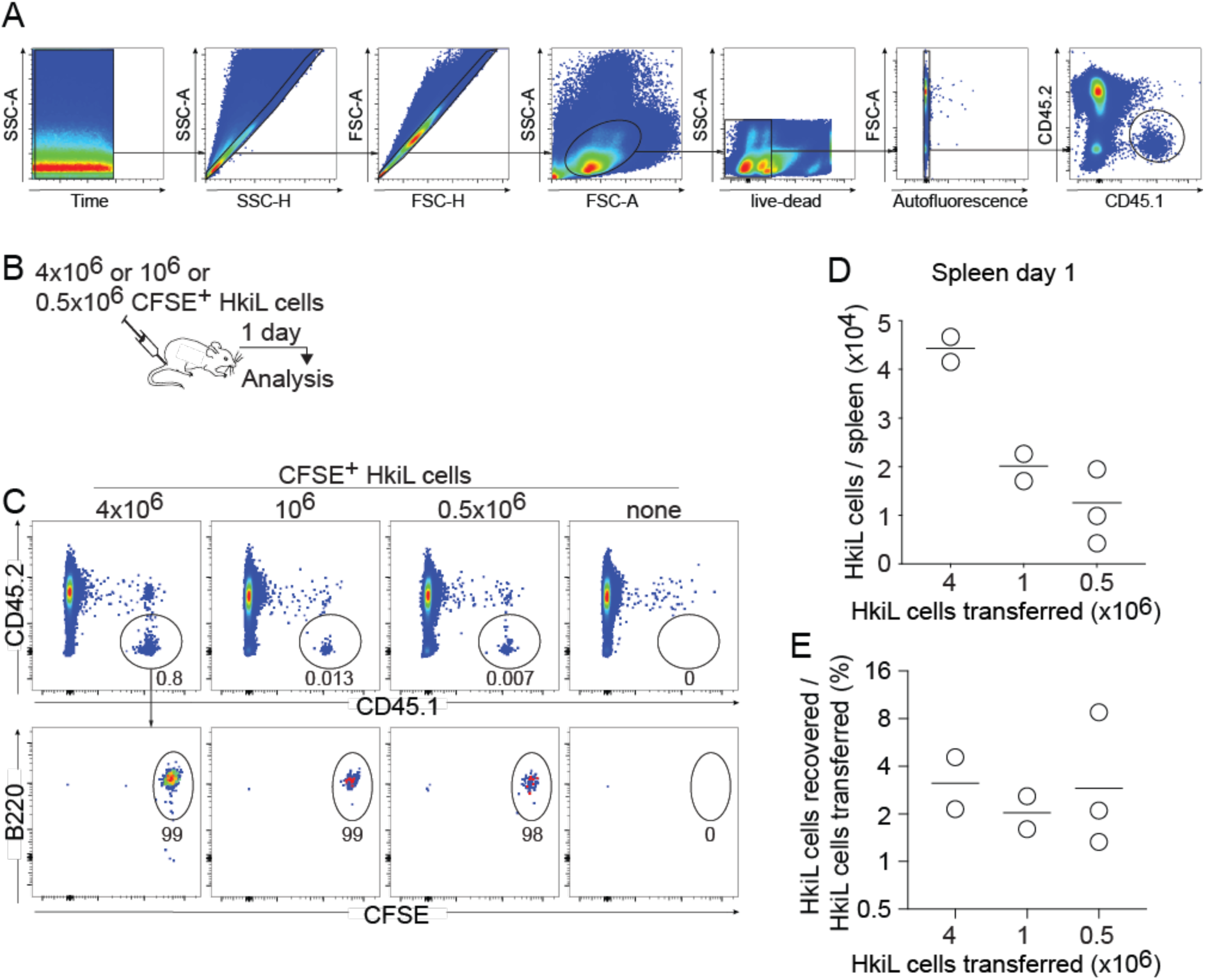
FACS gating strategy for the analysis of adoptively transferred HkiL cells and their splenic take. A: Gating strategy for the analysis of adoptively transferred HkiL cells. B-E: We transferred either 4×10^6^, 10^6^ or or 5×10^5^ CFSE-labelled HkiL cells on d0 and counted their numbers in spleen 24 hours later (B). Representative FACS plots (C; 2 mice receiving 4×10^6^, 2 mice receiving 10^6^ cells and 3 mice receiving 5×10^5^ HkiL cells) pre-gated on live lymphocytes (top) as shown in (A), with the gated HkiL cells analyzed for expression of B220 and CFSE (bottom), documenting homogenously high CFSE levels and uniform B220 expression, as expected for naïve B cells. Total live HkiL cells (D) and the splenic take (E) calculated as (number of HkiL cells recovered from spleen / number of HkiL cells transferred). Numbers in FACS plots indicate the percentage of gated cells as the group mean (C). Symbols in D and E show individual mice with the mean indicated.

**Figure S2:**
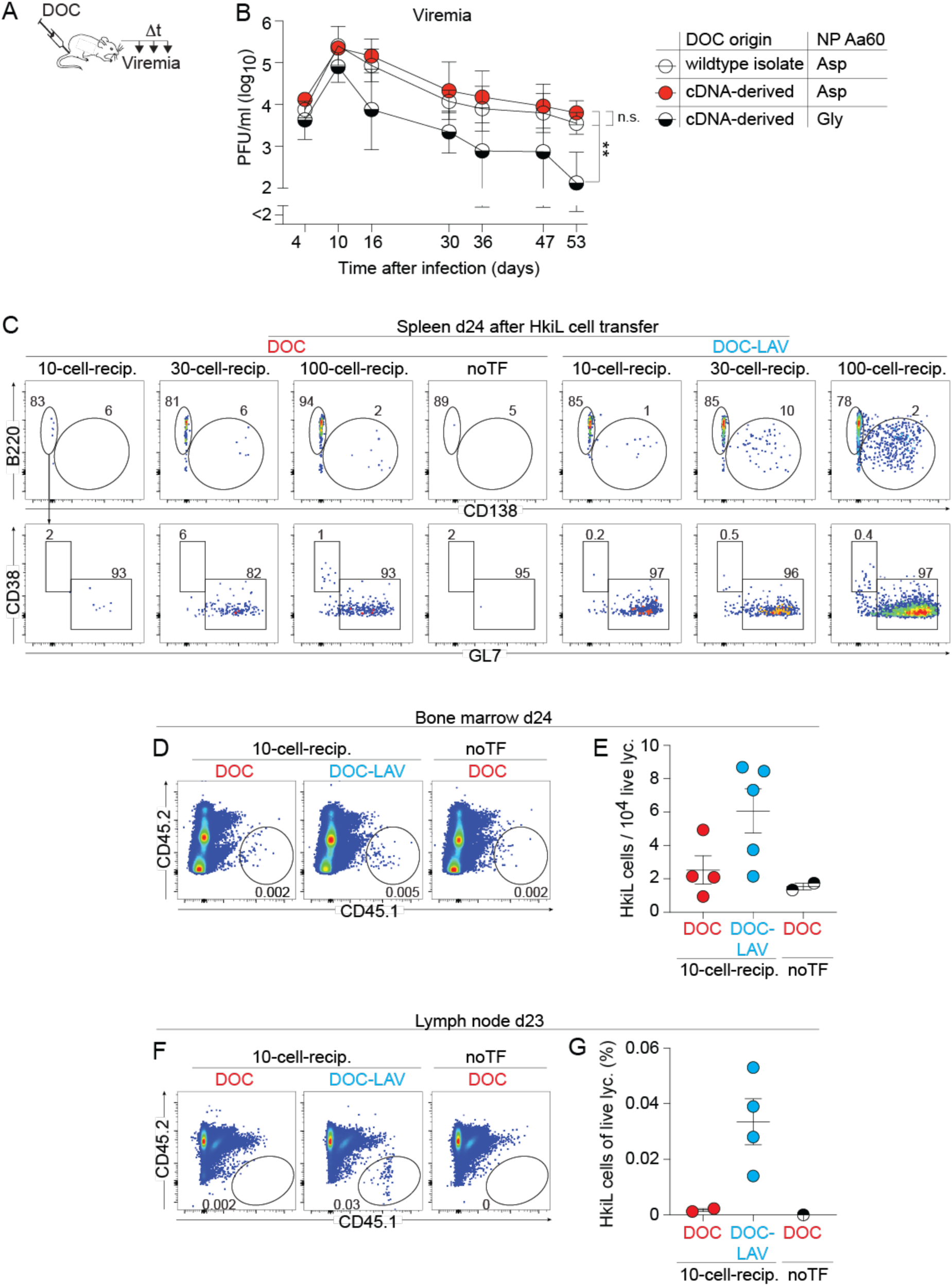
DOC reverse genetic system, differentiation phenotype of HkiL cells undergoing attrition and manifestation of attrition in lymph nodes and bone marrow. A-B: Based on cDNAs previously published^92^ and following validated techniques^96,124^ we established a reverse genetic system for DOC. We noticed, however, that when administered to mice, the cDNA-derived virus did not establish viremia at levels and of a durability comparable to our wildtype isolate DOC. We therefore revisited the viruses’ sequence and identified a coding difference (G60N) in the viral nucleoprotein (NP). Hence, we reverted Aa60 in the NP ORF of our cDNA rescue system to asparagine and generated the corresponding cDNA-derived DOC virus. We then infected mice with either one of three versions of DOC (A,B): wildtype isolate DOC, cDNA-derived DOC with NP Aa60 Asp or cDNA-derived DOC with NP Aa60 Gly. We collected blood over time and determined viremia (B). Reverting NP Aa60 in the cDNA-derived DOC from Gly to Asp restored its ability to establish viremia at levels and with a durability indistinguishable from the wildtype DOC isolate. The DOC and DOC-LAV viruses used in the remainder study have Asp at Aa60 of NP. C-E: In the experiment to Fig. 2 we infected mice with DOC or DOC-LAV on d-6. On d0 we engrafted either 10, 30 or 100 HkiL cells or none (noTF) and analyzed their progeny in spleen on d24 (C). HkiL cells gated as shown in Fig. 2B were analyzed for B220 and CD138 expression (top), and the B220^+^CD138^-^ B cell subset was further analyzed for GL7^+^CD38^-^ GC B cells and GL7^-^CD38^+^ memory B cells (bottom). From the 10-cell-recipient groups we further enumerated HkiL cell progeny in the bone marrow (D,E). Representative FACS plots are shown (D) and the relative abundance in bone marrow (E) was analyzed. F,G: In an experiment conducted analogously to the one in (C-E) we analyzed HkiL progeny in inguinal lymph nodes on d23 after transferring 10 HkiL cells. Representative FACS plots (F; n=2 in DOC, n=4 in DOC-LAV-infected groups, n=1 in noTf group), pre-gated on live lymphocytes (Fig. S1A) and the relative abundance of HkiL cells (G) are shown. Numbers in FACS plots indicate the percentage of gated cells as the group mean. Symbols in (B) show the mean±SEM of 3-5 mice per group. Symbols in E,G show individual mice with the mean±SEM indicated. Viremia on d53 in (B) was compared by performing one-way ANOVA with Tukey’s post test. Data in A-G show one representative of two similar experiments. **: p<0.01; n.s.: not statistically significant.

**Figure S3:**
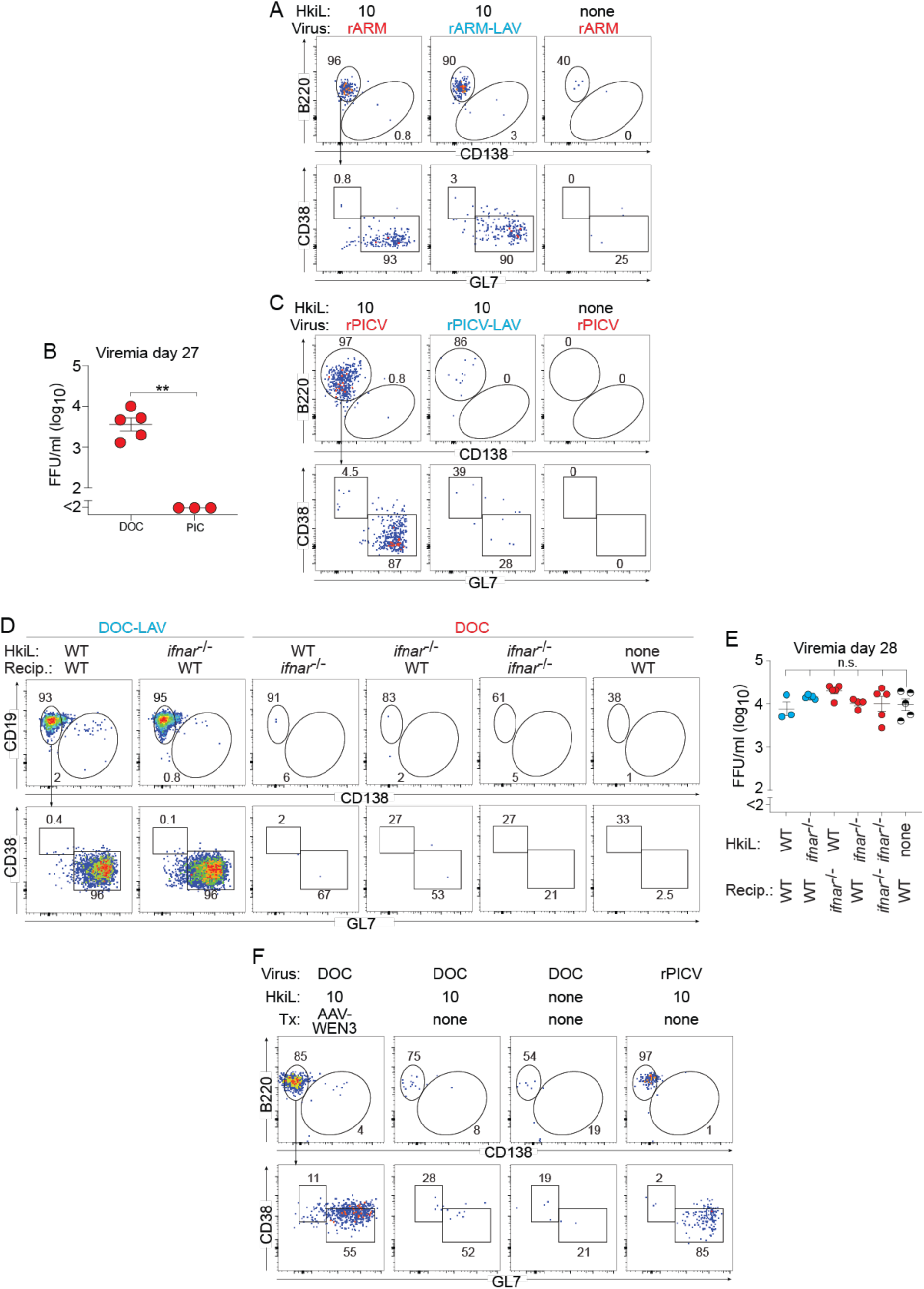
Differentiation of HkiL cells responding to rARM and rPICV, failure of rPICV to establish viremia, and the impact of IFN-I signaling as well as of AAV-WEN3 therapy on HkiL cell differentiation. A: In the experiment to Fig. 3A-C we infected mice with rARM or rARM-LAV on d-6 and on d0 engrafted 10 HkiL cells or none (noTf). HkiL progeny in spleen were analyzed on d20. Representative FACS plots (n=4 mice except n=2 mice in noTf group) of HkiL cells as gated in Fig. 3B, analyzed for B220 and CD138 expression (top). The B220^+^CD138^-^ B cell subset was further analyzed for GL7^+^CD38^-^ GC B cells and GL7^-^CD38^+^ memory B cells (bottom). B: We infected mice with DOC or rPICV, engrafted 10 cells at d0 and analyzed viremia on d27. Symbols show individual mice with the mean±SEM indicated. C: In the experiment to Fig. 3D-F we infected mice with rPICV or rPICV-LAV on d-6 and on d0 engrafted 10 HkiL cells or none (noTF). HkiL progeny in spleen were analyzed on d27. Representative FACS plots (C; n=4 mice for rPIC, n=3 mice for rPICV-LAV group, n=1 for noTF) of HkiL cells as gated in Fig. 3E, analyzed for B220 and CD138 expression (top). The B220^+^CD138^-^ B cell subset was further analyzed for GL7^+^CD38^-^ GC B cells and GL7^-^CD38^+^ memory B cells (bottom). D,E: In the experiment to Fig. 3G-I we infected *ifnar*^-/-^ and WT control recipients with DOC or DOC-LAV on d-6. On d0 we engrafted them with either 10 HkiL cells or 10 HkiL-*ifnar*^-/-^ cells or left them without cell transfer (”none”, noTF) in the combinations indicated. HkiL and HkiL-*ifnar*^-/-^ progeny (CD45.1^+^) were analyzed on d28. Representative FACS plots (n=5 except HkiL into DOC-LAV-infected WT mice (n=3) and HkiL-*ifnar*^-/-^ into DOC-infected wt mice (n=4)) of the cells gated in Fig. 3H are shown in (D), analyzed for CD19 and CD138 expression (top). The CD19^+^CD138^-^ B cell subset was further analyzed for GL7^+^CD38^-^ GC B cells and GL7^-^CD38^+^ memory B cells (bottom). Viremia was determined on d28 (E). F: In the experiment to Fig. 3J-M we infected mice with DOC or rPICV on d-6, and on d0 engrafted 10 HkiL cells or none (noTF) as indicated. On d5 we administered AAV-WEN3 to one DOC-infected group. HkiL progeny in spleen were analyzed on d24. Representative FACS plots (n=9 AAV-WEN3-treated HkiL recipients, n=8 recipients without AAV-WEN3, n=9 rPICV-immunized HkiL recipients, n=5 noTF controls) of HkiL cells as gated in Fig. 3K are shown, analyzed for B220 and CD138 expression (top). The B220^+^CD138^-^ B cell subset was further analyzed for GL7^+^CD38^-^ GC B cells and GL7^-^CD38^+^ memory B cells (bottom). We performed unpaired Student’s t-test for statistical analysis of the values in (B), one-way ANOVA with Dunnett’s post-test was used in (E) to compare viremia of the various HkiL recipients against the noTf group. Data in (A-E) show one out of two similar experiments, (F) reports results from two combined experiments. **: p<0.01; n.s.: not statistically significant.

**Figure S4:**
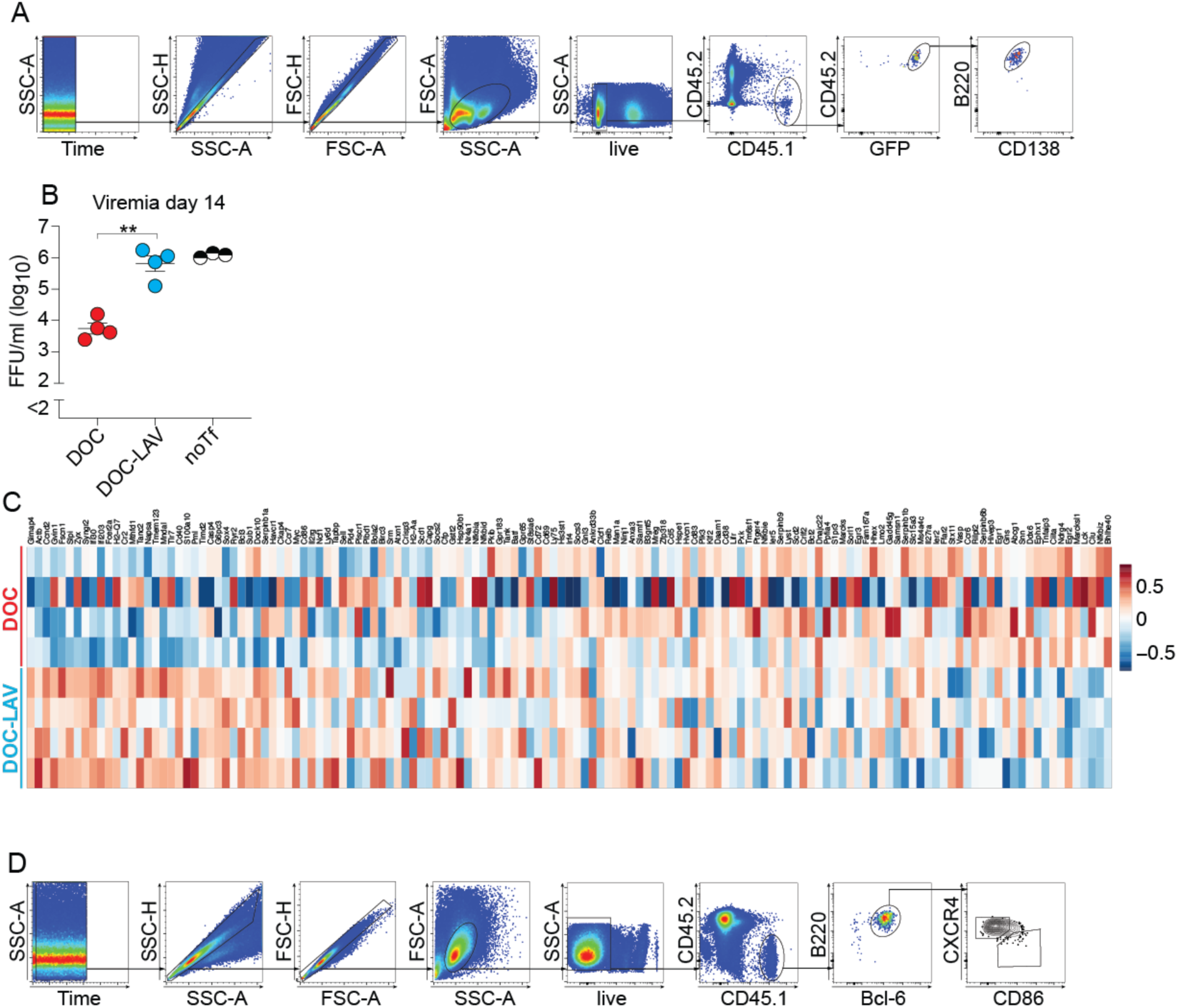
FACS gating strategy for the analysis of adoptively transferred HkiL cells, viremia in DOC- and DOC-LAV-infected HkiL cell recipients, and these cells’ expression of GC light zone signature genes. A: Gating strategy for the sorting of HkiL-GFP cells in the experiment reported on in Figs. 4F-H and 5A-D. B: Viral titers in the blood of the animals from which HkiL cells were sorted for the analyses reported in Figs. 4F-H and 5A-D. An additional group of DOC-infected mice without HkiL transfer (noTf; not reported on in Figs. 4 or 5) was included for reference. Symbols show individual mice with the mean±SEM indicated. Unpaired Student’s t test was performed for statistical analysis. **: p<0.01 C. Heatmaps as shown in Fig. 4G with individual gene names indicated. D: Gating strategy for the analysis of Bcl6^+^B220^+^ HkiL GC B cells in the experiment to Fig. 4I-L.

**Figure S5:**
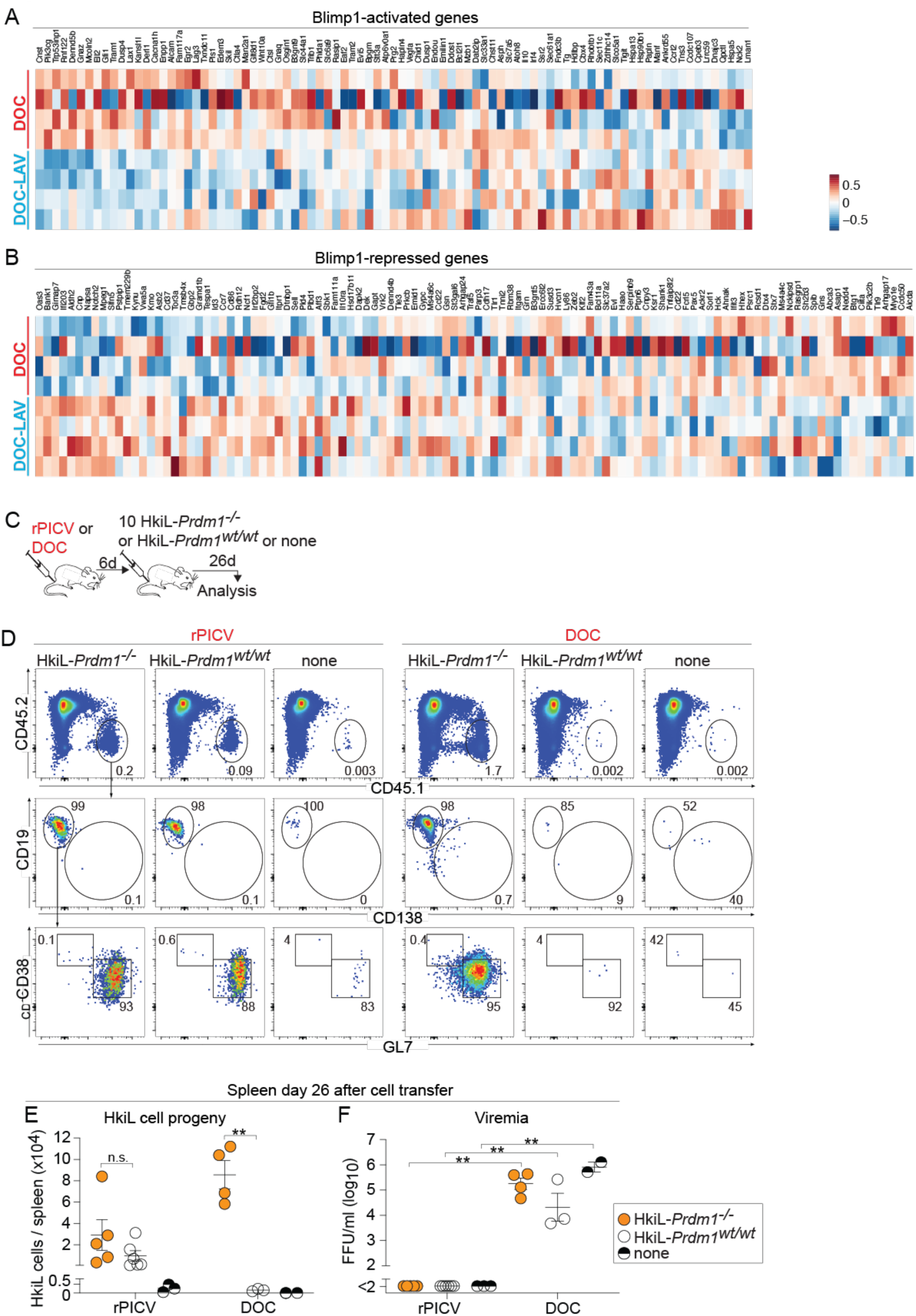
Differential expression of Blimp-1-regulated genes in HkiL cells responding to DOC and DOC-LAV, and comparable response of *Prdm1*-deficient and -sufficient HkiL cells to rPICV immunization. A,B: Heatmaps as shown in Fig. 5A,C with individual gene names indicated. C-F: We infected recipient mice with DOC or rPICV on d-6, and on d0 we engrafted either 10 HkiL-*Prdm1*^-/-^ or 10 HkiL-*Prdm1*^wt/wt^ cells or none (noTF). These cells’ progeny in spleen were analyzed on d26 (C). Representative FACS plots (D; n=5 rPICV-immunized HkiL-*Prdm1*^-/-^ cell recipients, n=6 rPICV-immunized HkiL-*Prdm1*^wt/wt^ cell recipients, n=3 rPICV-infected noTf mice, n=4 DOC-infected HkiL-*Prdm1*^-/-^ cell recipients, n=3 DOC-infected HkiL-*Prdm1*^wt/wt^ cell recipients, n=2 DOC-infected noTf mice), pre-gated on live lymphocytes (Fig. S1A). HkiL progeny (CD45.1^+^, top) were analyzed for expression of CD19 and CD138 (center), and their CD19^+^CD138^—^ subset was analyzed for GL7 and CD38 expression (bottom). HkiL cell progeny numbers (E) and viremia (F) were determined on d26, too. Numbers in FACS plots indicate the percentage of gated cells as the group’s mean. Pair-wise comparisons in (E,F) were performed by unpaired Student’s t tests with Bonferroni correction. Results in (D-F) show one representative out of two similar experiments. **: p<0.01; n.s.: not statistically significant.

**Table SI.**
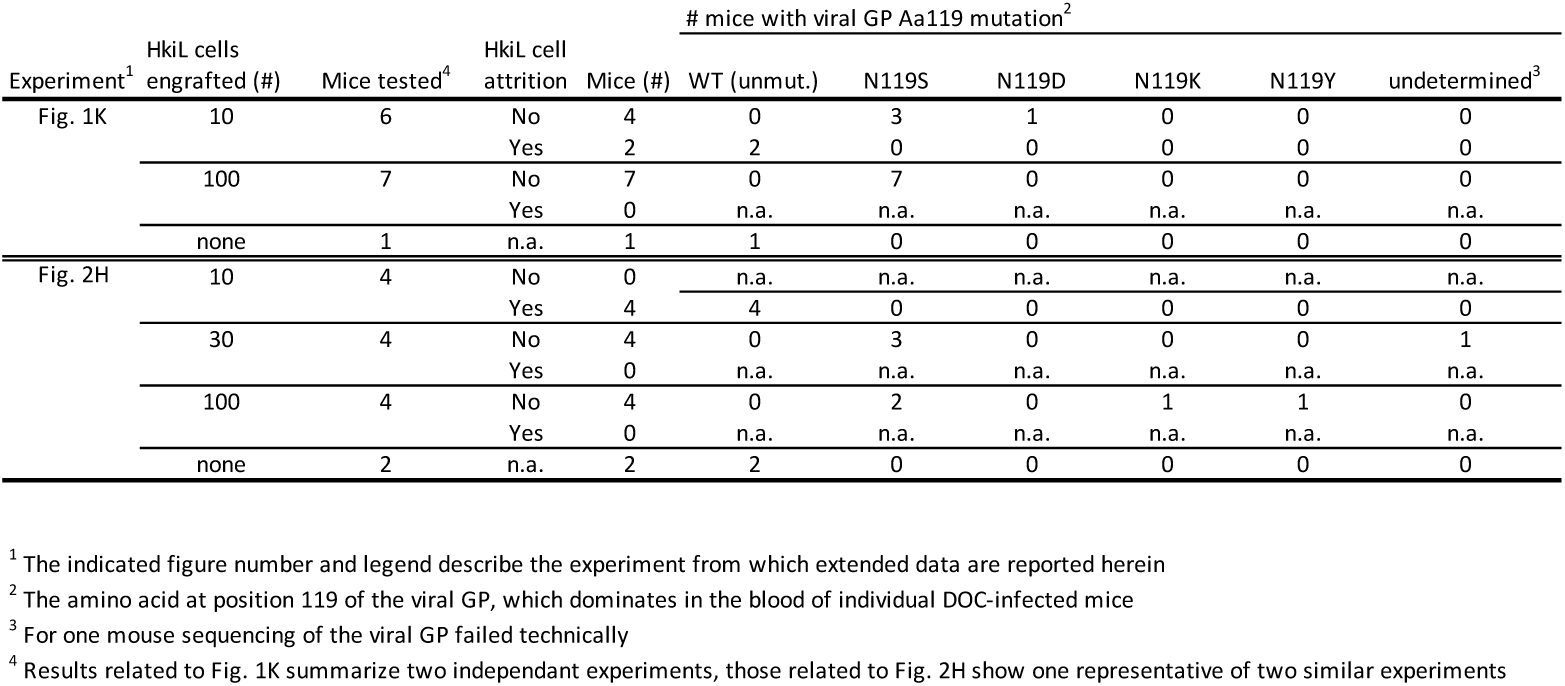

